# Dispatched conformational dynamics couples transmembrane Na^+^ flux to release of lipid-modified Hedgehog signal

**DOI:** 10.1101/2020.09.05.284026

**Authors:** Qianqian Wang, Daniel E. Asarnow, Ke Ding, Yunxiao Zhang, Yong Ma, Yifan Cheng, Philip A. Beachy

**Affiliations:** Institute for Stem Cell Biology and Regenerative Medicine, Stanford University School of Medicine, Stanford, California 94305, USA; Department of Biochemistry and Biophysics, University of California, San Francisco, California 94158, USA; Department of Molecular Biology and Genetics, Johns Hopkins University School of Medicine, Baltimore, CA 21205; Howard Hughes Medical Institute, University of California, San Francisco, California 94158, USA; Departments of Urology, and Developmental Biology, Stanford University School of Medicine, Stanford, California 94305, USA

**Author notes:** Suzhou Institute of Biomedical Engineering and Technology, Chinese Academy of Sciences, 88 Keling Road, SND, Suzhou, Jiangsu 215163, China. Howard Hughes Medical Institute, Neuroscience Department, The Scripps Research Institute, La Jolla, CA 92037, USA. These authors contributed equally to this work (Qianqian Wang, Daniel E. Asarnow). Corresponding author. (Y.C.), (P.A.B.).

## Abstract

The DISP1 protein, related to the NPC1 and PTCH1 cholesterol transporters and to H^+^-driven transporters of the RND family, enables tissue patterning activity of the lipid-modified Hedgehog protein by releasing it from tightly-localized sites of embryonic expression. A 2.5 Å resolution cryo-EM structure of DISP1 revealed three Na^+^ ions coordinated within a channel that traverses its transmembrane domain. This channel and Na^+^-binding are disrupted in DISP1-NNN (3.4 Å resolution), a variant with isosteric substitutions for three intramembrane aspartates that each coordinate and neutralize charge for one of the three Na^+^ ions. DISP1-NNN and other variants that singly disrupt the Na^+^ sites retain binding to the lipid-modified Hedgehog protein, but most fail to export Hedgehog. The Sonic hedgehog signal (ShhN) interacts with the DISP1 extracellular domains (2.7 Å complex structure), including an unusual extended DISP1 arm, at a location well above the membrane. Variability analysis reveals a dynamic series of DISP1 conformations, only a restricted subset of which appear to bind ShhN. The bound and unbound DISP1 conformations display distinct Na^+^ site occupancies; these differences, in conjunction with the unusual locations of certain lipids bound to DISP1, suggest a mechanism by which transmembrane Na^+^ flux may power extraction of the lipid-linked Hedgehog signal from the membrane, thus enabling its tissue patterning activity. The Na^+^-coordinating residues resolved in our DISP1 structures are wholly or partly conserved in some of the other metazoan RND family members, such as PTCH1 and NPC1, suggesting the utilization of Na^+^ flux to power their conformationally-driven activities.

## Introduction

During embryogenesis the Hedgehog protein signal projects its tissue patterning influence many cells beyond its sites of expression in “organizers” such as the vertebrate notochord, the floor plate of the neural tube, and the zone of polarizing activity in developing limb[1–4]. The mature Hedgehog protein signal (ShhNp) is covalently lipid-modified[5] by autoprocessing-mediated cholesterol attachment at its carboxy-terminus[6] and acyl transferase-mediated attachment of palmitate at its amino-terminus[7,8]. The resulting dually lipid-modified protein is tightly membrane associated and thus requires dedicated machinery for its packaging and release in long-range patterning. This machinery includes the DISP1 protein[9–13], required in Hedgehog-producing cells, and SCUBE, a secreted protein that can be supplied by other cells that do not produce or respond to the Hedgehog signal[14–16]. DISP1 is related to the PTCH1 and NPC1 cholesterol transporters [17,18], and to the large prokaryotic family of RND transporters that export substrates from the periplasm by a proton-driven antiport mechanism[19,20]. Curiously, although the DISP1 protein and the Hedgehog receptor PTCH1 are similar in sequence and transmembrane topology and both interact with the lipid-modified Hh signal, their activities in Hh signaling are strikingly different (Fig. S1A). DISP1 promotes pathway activation by releasing Hh from producing cells[9–13]: its embryonic mutant phenotype in mice is more severe than that of the *Sonic hedgehog* (*Shh*) or *Indian hedgehog* (*Ihh*) single mutants and resembles that of the *Shh;Ihh* double mutant, and of a homozygous knockout of the essential Hedgehog response component *Smoothened* (*Smo*)[10,11,21]. The PTCH1 protein in contrast functions to attenuate pathway activity by inhibiting SMO, and *Ptch1* homozygotes show widespread ectopic pathway activity[22]. The relationship between these opposing functions at distinct pathway levels is one of genetic epistasis, in which the *Ptch1^−/−^* phenotype, widespread pathway activation, prevails in *Ptch1^−/−^;Disp1^−/−^* doubly homozygous mutant embryos[10,13] (Fig. S1B).

## Results

### DISP1 structure determination

To illuminate the mechanism of DISP1 action and its mechanistic differences from PTCH1 we set out to determine the structure of DISP1. As murine full-length DISP1 protein was poorly expressed in HEK293 cells (Fig. S1C,D) we tested a variety of constructs, and ultimately settled on an N-terminal truncation, hereafter referred to as DISP1-A (Fig. S2), as an alteration that effectively stabilizes the protein and enhances its expression (Fig. S1C,D), thus rendering it suitable for purification and structure determination. DISP1-A mediated the release of ShhNp (autoprocessed, lipid-modified Shh protein) into cell culture medium containing SCUBE2 upon transfection into *Disp^−/−^* mouse embryonic fibroblasts[10,15] (MEFs), indicating preservation of function in DISP1-A (Fig. S1E).

Following detergent solubilization, DISP1-A was purified into a mixture of LMNG/GDN/CHS, and a size exclusion chromatography fraction corresponding to monomeric DISP1-A was found to be efficiently cleaved at a position consistent with a recently reported essential furin site[23] (Fig. S1F). This sample proved amenable to single-particle cryo-EM, and 3D classification within a single dataset revealed high-resolution structures of two major conformations (Fig. S3, S4; Table S1). We present here the basic features of a model of DISP1-A built from the density of one of these two classes, referred to as R (relaxed, 2.5 Å resolution; Fig. 1A,B); the differences between R and the other major class T (tense, 2.5 Å resolution) are described in Fig. S5).

**Fig. 1.**
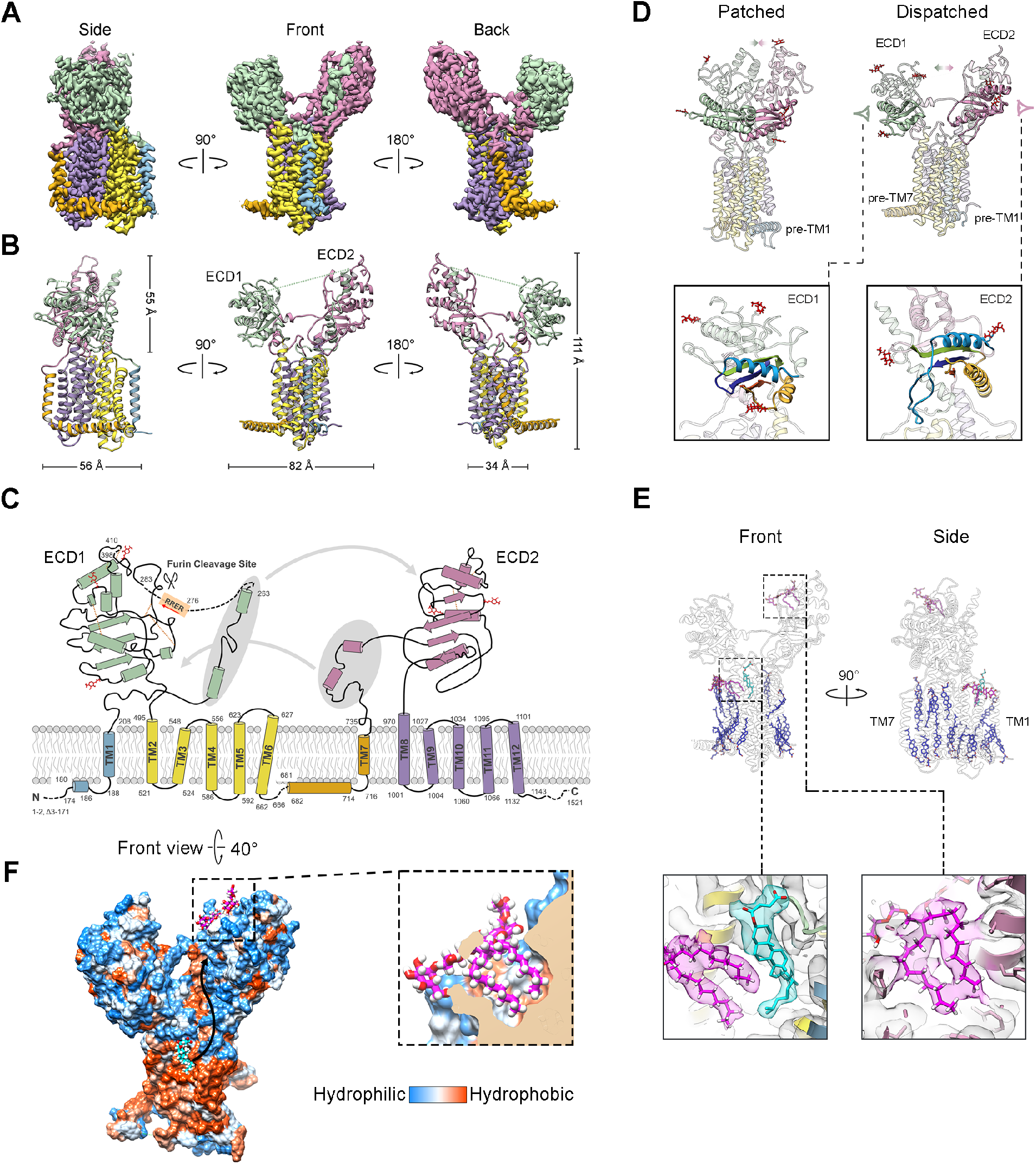
Structure, conserved ECD features, and lipid binding sites on DISP1. **A-B**, Corresponding views of the cryo-EM map (**A**) and atomic model (**B**) of DISP1-A in R conformation (see Fig. S3–5). Domains are colored as follows: pre-TM1 helix and TM1, malachite; ECD1, lime; TM2–TM7, yellow; pre-TM7 and TM7, orange; ECD2, pink; TM8–TM12, lavender. **C**, Topology diagram of DISP1-A showing secondary structure elements (same colors as in **A-B**), disulfide bonds (gold dashed lines), glycosylation sites (red symbols) and unresolved regions (black dashed lines). Secondary elements exchanged between extracellular loops to form ECD1 and ECD2 are highlighted by oval shadows. **D**, Structural comparison between DISP1-A and mouse PTCH1. The ECDs, closely apposed in PTCH1 to form a conduit for sterol transport, in DISP1-A are splayed apart. Conserved ferredoxin-like α+β open-faced sandwich folds are highlighted (lime for ECD1 and pink for ECD2). Magnified views in spectral sequence from N to C termini of the sandwich folds in DISP1-A ECD1 (bottom left) and ECD2 (bottom right). Red symbols indicate glycosylation sites. **E**, Front and side views of DISP1-A (pale ribbon) with resolved LMNG (magenta) and CHS (blue) molecules. A front-side lifted sterol molecule (cyan) is located at a location corresponding to the sterol sensing domain (SSD). Atomic model (stick) and corresponding density map showing a magnified view (bottom left) of the front-side lifted sterol (cyan), which associates with a tilted LMNG molecule (magenta). Similar magnified view (bottom right) of a hydrophobic cavity in the distal portion of ECD2, far from the transmembrane region. This cavity is occupied by another LMNG molecule (magenta). **F**, Representation of surface hydrophobicity, viewed from a top-front position of DISP1 (left) and a close-up cut-away view of the ECD2 hydrophobic cavity (right). A hydrophobic track beginning near the front-side lifted sterol (cyan) extends to the proximal edge of the ECD2 hydrophobic cavity.

The DISP1-A structure resembles that of PTCH1, with twelve transmembrane (TM) helices derived from apparent duplication of a six-TM unit, with each six-TM unit (TM1-6, TM7-12) preceded by an intracellular transverse helix (pre-TM1 and pre-TM7) and containing large extracellular loops deriving from corresponding positions between TM1 and TM2 and between TM7 and TM8 (Fig. 1C). Unlike PTCH1, in which the extracellular domains (ECDs) are juxtaposed to form a conduit for sterol transport[17], in DISP1 the ECDs are splayed apart in a Y-shaped manner from their points of origin in the TM domain (Fig. 1A,B,D). Although each ECD is predominantly constructed from one of the extracellular loops, each also accepts several secondary structure elements from the other loop, creating linkers that enclose a bowl-shaped space between the two domains. One of these linkers, within the TM1-TM2 loop, is the site of furin cleavage (RERR, 276-279)[23], which opens the wall of the bowl at a location we designate as the “front”. The ECDs contain 7 disulfide bonds between Cys pairs, 5 in ECD1 and 2 in ECD2 (Fig. 1C, Fig. S2), with clear densities visible for all but one of the corresponding bonds between side-chain sulfurs (Fig. S6). In addition, 5 N-linked glycosylation sites, 3 in ECD1 (N362, N390 and N475) and 2 in ECD2 (N834 and N915), can be inferred from additional densities that extend from N-X-S/T sequences in the extracellular loops (Fig. 1C,D). With the exception of pre-TM1 and pre-TM7, most of the intracellular sequence in DISP1-A is not resolved and not modeled, including four amino-terminal residues (1-2, 172-3), 14 residues preceding pre-TM7 (667-680), and 378 residues at the carboxy-terminus (1144-1521). In the extracellular portion, 19 residues that include the furin cleavage site (264-282) and 11 residues in ECD1 (399-409) are not sufficiently well-resolved to support modeling.

The relationship of DISP1 to PTCH1 is clear from their closely similar TM domains, but is also evident from the presence of similar ferredoxin-like α+β open-faced sandwich folds in the membrane-proximal parts of ECD1 and ECD2 of both proteins (Fig. 1D). More distal structures within ECD1 and ECD2 derive from insertions into the peripheral loops of the ferredoxin-like folds and appear to be structurally unrelated to each other or to distal ECD structures in PTCH1.

The high quality of the DISP1-A structure allowed us to confidently model 20 lipid or detergent molecules, most of them located at annular positions between the TM domain and the boundary of the detergent micelle. Eighteen of these were modeled as CHS, eleven in positions corresponding to the inner leaflet of the membrane and seven to the outer leaflet. These molecules are positioned with polar groups oriented toward the inner and outer surfaces of the membrane, presenting the appearance of a sterol bilayer (Fig. 1E). Of the seven outer leaflet sterols, two are unusual in their raised positions relative to the membrane, at corresponding positions on the front and back sides of DISP1-A. The front-side lifted sterol is located in a pocket bounded by TMs 1 and 2 on the periphery, TM3 from the bottom, and TM4 internally, and the back-side lifted sterol between TMs 7 and 8 peripherally, TM9 from the bottom, and TM10 internally. The front-side lifted sterol is particularly interesting because it appears to protrude outward from its hydrophobic surroundings and because a nearby lipidic group, modeled as LMNG with its acyl chains oriented somewhat laterally, appears to lever the lifted sterol upward (Fig. 1E). One other notable feature, located at the distal tip of ECD2, is a hydrophobic cavity filled with non-proteogenic density, which we modeled as accommodating the acyl chains of LMNG (Fig. 1F).

### Na^+^ ions in a transmembrane tunnel

A close inspection of the TM domains revealed three striking non-proteogenic intramembrane densities (sites I, II and III in Fig. 2A), each associated with one of the three acidic residues located in TM4 and TM10 that are conserved in PTCH1 (Fig. 2B). We interpret these densities as bound metal cations, which appear well-coordinated by oxygens from side-chain hydroxyls and main-chain carbonyls, and at least one oxygen from water (site III) (Fig. 2C). The coordination number of the bound ions is five for sites I and II and six for site III, and the center-to-center distances between ions and their associated oxygens average 2.52 Å, 2.54 Å, and 2.52 Å at sites I, II, and III, respectively. The association of each density with just a single carboxylate and with oxygens from hydroxyls and main chain carbonyls rules out alkaline earth metal ions such as Ca^++^ or Mg^++^, which typically require additional charge neutralization, display a more rigid geometry and a tighter coordination sphere, and are rarely coordinated by main-chain carbonyls. In addition, the average bond distances match those characteristic of Na^+^ (2.4 – 2.5 Å) rather than K^+^ (2.7-3.2 Å)[24]. On the basis of geometry (Na^+^ can be coordinated by five or six ligands), bond lengths, and degree of available charge neutralization we conclude that the densities at sites I, II, and III derive from Na^+^ [24]. Assessment of our coordination site models with the software package *CheckMyMetal*[24] web server further validates Na^+^ as the most likely ions bound at these sites. The assignment of these densities as Na^+^ is also supported by the tight conservation of intramembrane acidic amino acids and other coordinating ligands between DISP1 and PTCH1 (Fig. 2D), the latter of which has been shown to depend on Na^+^ for its function in regulating SMO[25].

**Fig. 2.**
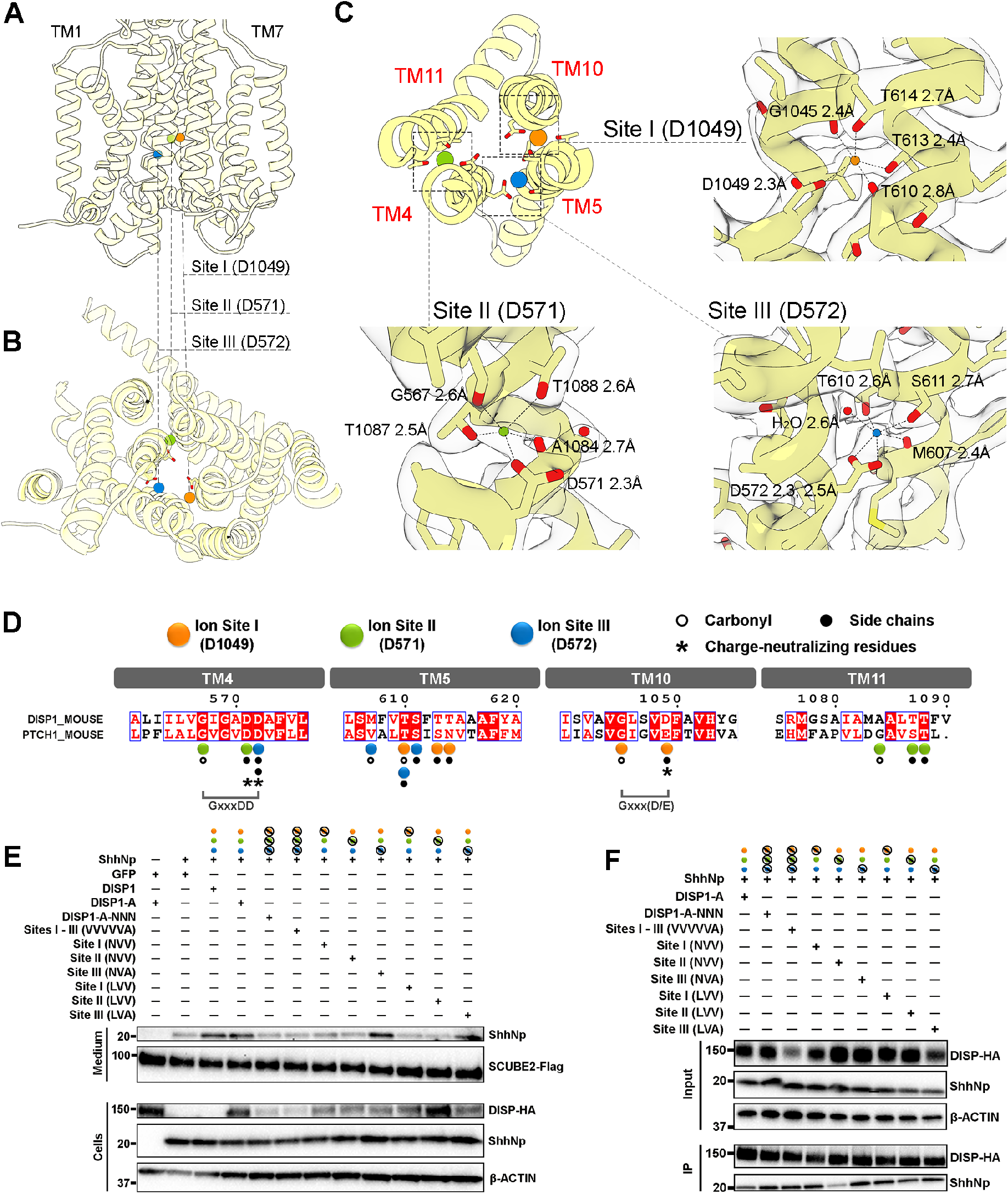
Three Na^+^ densities in the transmembrane domain. **A-B**, Side (**A**) and extracellular (**B**) views of three Na^+^ ion binding sites within the transmembrane domain, each labeled with its associated intra-membrane Asp. **C**, Close-up views of the full set of coordinating ligands for each of the three Na^+^ ions is shown with protein-derived densities superimposed upon the atomic model of DISP1-A (overview, top left). Dashed lines denote coordination bonds and corresponding center-to-center distances between Na^+^ ions and associated oxygens. Contour levels are 8σ for sites I and II, and 5σ for site III. **D**, Na^+^-interacting residues in DISP1 are conserved in PTCH1 (see also Fig. S10). **E**, Culture media and cell lysates from transiently transfected Disp−/− MEFs were probed by immunoblotting for DISP-HA, ShhNp, and SCUBE2. ShhNp released into culture medium from cells expressing DISP1-A-NNN, DISP1-A-VVVVVA, DISP1-A-NVV (Site I), DISP1-A-NVV (Site II), DISP1-A-LVV (Site I), and DISP1-A-LVV (Site II) is similar to the GFP control, indicating lack of export activity. β-ACTIN, loading control. **F**, DISP-ShhNp binding assay using HEK293 with stably integrated constructs for doxycycline-inducible expression of full-length Shh[15]. DISP1-A variants tagged with SBP were immunoprecipitated with Streptavidin resin, and ShhNp detected by Western blot. Alterations in ion site residues as follows: DISP1-A-NNN, D571N/D572N/D1049N; DISP1-A-VVVVVA, T613V/T614V/T1087V/T1088V/T610V/S611A; DISP1-A-NVV (Site I), D1049N/T613V/T614V; DISP1-A-NVV (Site II), D571N/T1087V/T1088V; DISP1-A-NVA (Site III), D572N/T610V/S611A; DISP1-A-LVV (Site I), D1049L/T613V/T614V; DISP1-A-LVV (Site II), D571L/T1087V/T1088V; DISP1-A-LVA (Site III), D572L/T610V/S611A.

DISP1 function *in vivo* has long been known[10] to be impaired by simultaneous alteration of D571, D572, and D1049, the acidic residues within TM4 and TM10 that we find here provide charge neutralization and coordinate the three intramembrane Na^+^ ions in DISP1-A (Fig. 2A-C). We confirmed that substitution of these residues by asparagine in the context of our DISP1-A construct (DISP1-A-NNN) also disrupts ShhNp release in our *in vitro* assay (Fig. 2E). We also noted, however, that asparagine substitution for individual charge-neutralizing aspartates does not inactivate DISP1-A (not shown). Reasoning that alteration of an individual aspartate may not suffice to completely disrupt Na^+^ binding, we subsequently altered other residues at each site that supply coordinating side-chain oxygens. We found that disruption of individual Na^+^ sites I or II, but not site III, abrogate ShhNp release in our *in vitro* assay (Fig. 2E). In the mutants that exclusively disrupt site I or site II, the requirement for alteration of the entire set of coordinating side-chains suggests joint contributions of all coordinating residues at any individual site in mediating Na^+^ binding. We also left intact the aspartates, while altering residues whose side chains provide coordinating oxygens, simultaneously at all three ion sites (site I: T613 and T614; site II: T1087 and T1088; site III: T610 and S611). Indeed, this protein (DISP1-A-VVVVVA) fails to release ShhNp (Fig. 2E), again highlighting the importance of the non-aspartate, Na^+^-coordinating residues. As all of these variant proteins, like DISP1A-NNN, are stable and retain at the ability to bind ShhNp (Fig. 2F), we conclude that they are well-folded and that defects in release of the ShhNp protein result from their inability to utilize Na^+^ flux in a productive activity cycle (see below).

Purification of DISP1-A-NNN and structure determination by cryo-EM at 3.4 Å resolution (Fig. S7A; Table S1) revealed that the Na^+^ ion densities at sites I, II, and III in DISP1-A are indeed absent (Fig. 3A), and that some of the liganding side chains are rearranged, with rotamers no longer in Na^+^-coordinating position (Fig. 3A). One other striking difference is that the hydrophobic cavity in distal ECD2, although present in the DISP1-A-NNN structure, is distinctly shaped and does not contain the LMNG density present in DISP1-A (Fig. S7B).

**Fig. 3.**
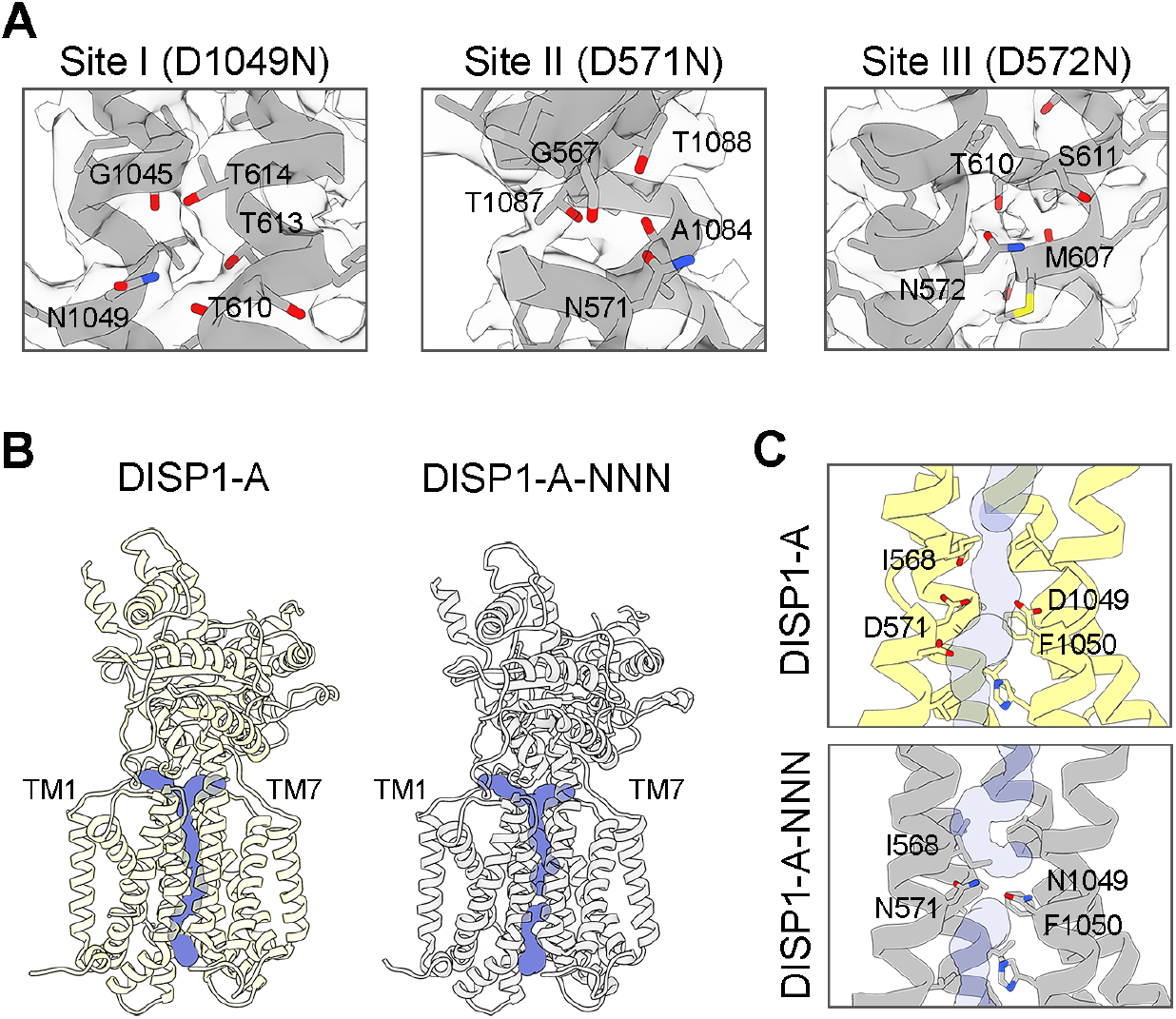
Disrupted ion conductance in the structure of DISP1-A-NNN. **A**, Close-up views of the three Na^+^ ion sites in DISP1-A-NNN show no discernible Na^+^ densities, even at 2σ level (Na^+^ densities in DISP1-A were clearly seen at 8σ for sites I and II or 5σ for site III). **B**, Tunnels emanating from the Na^+^ coordination nexus in DISP1-A, R conformation (khaki), and DISP1-A-NNN (grey) were computed using the CAVER 3 PyMOL plugin with a probe of radius 1.0 Å. The conduction pathway is occluded in DISP1-A-NNN. **C**, A close-up view of CAVER tunnels and the ion coordination nexus shows that in DISP1-A-NNN, the ion conduction pathway is blocked by I568, N571, N1049 and F1050. The intracellular exit is blocked by N571, N1049 and F1050.

The coincident loss of intramembrane Na^+^ and function in DISP1-A-NNN suggests that chemiosmotic force derived from high extracellular Na^+^ powers DISP1 action in exporting lipid-modified Hedgehog protein. Exploitation of this Na^+^ gradient, however, implies Na^+^ flux across the membrane. Using the Caver algorithm[26], we found a continuous channel within the DISP transmembrane domain of appropriate diameter to permit Na^+^ flux across the membrane (Fig. 3B). Strikingly, however, this channel is disrupted in DISP1-A-NNN (Fig. 3B), with the side chain of Ile568, and those of Asn571 and Asn1049 (substituted for Asp571 and Asp1049), rotated to occlude the channel (Fig. 3C).

### Structure of a ShhN:DISP1-A complex

To gain further insight into the mechanism of DISP1 action, we determined its structure in complex with Hedgehog. As we have not yet found conditions that enable us to image DISP1 in complex with the lipid-modified Hedgehog protein (ShhNp), we imaged a DISP1-A preparation with truncated Shh protein containing residues 26-189 and lacking lipid modifications (ShhN) (Fig. S4, S8; Table S1). *Ab initio* reconstruction and refinement with cryoSPARC[27] yielded two major density classes, each with the typical Y-shaped appearance of DISP1-A, but with the largest class containing substantial additional density corresponding to ShhN perched near the top of the front side of DISP1-A, between ECD1 and ECD2 (Fig. 4A). Indeed, given the high quality of the map (2.7 Å overall resolution), a model of the of ShhN:DISP1-A complex was readily constructed (Fig. 4B). The ShhN portion of this map resolved residues 39-189 as well as the previously described Zn^++^ ion[28] and the di-atomic Ca^++^ cluster[29], along with all of the major secondary structure elements, and the expected positions of many side chains (Fig. 4A-B, Fig. S6, S8).

**Fig. 4.**
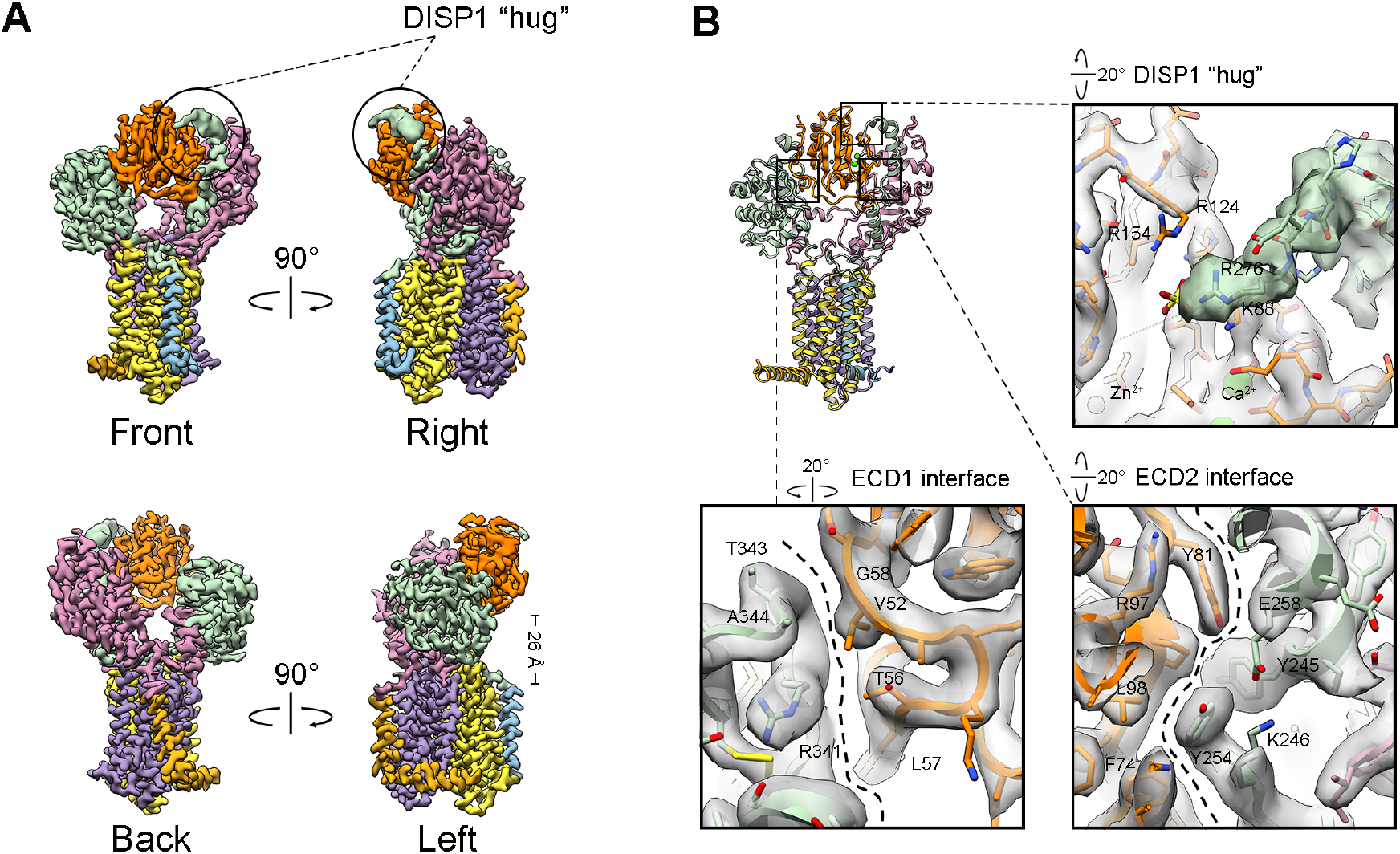
DISP1-A binds ShhNbetween its distal ECDs. Cryo-EM density map of a ShhN:DISP1-A complex at 2.7Å resolution (**A**) permits *de novo* modeling of ShhN at a location 26 Å above the TM domain of DISP1-A (**B)**. ShhN is in orange, DISP1-A colors are as in Fig. 1A-C. The furin-cleaved linker deriving from the TM1-TM2 loop, although not resolved in the DISP1-A apostructure, is stabilized in the complex to extend from ECD2 in a one-armed molecular hug of ShhN. The 989.3 Å^2^ of buried surface area between DISP1-A and ShhN derives mostly from three major interfaces that are shown in expanded form in (**B**). The ECD1 and ECD2 interfaces feature primarily van der Waals interactions, with the extensive ECD2 interface additionally including a hydrogen bond between ShhN K75 and the backbone carbonyl of DISP1-A R241 (located just below the expanded view; not shown). Near the tip of the furin-cleaved linker arm, R276 of DISP1-A forms an ionic interaction with a sulfate ion that also interacts with basic residues in ShhN (R124, R154, and K88).

The major interaction between DISP1-A and ShhN in the complex involves a pincer-like grasp of ShhN by ECD1 and ECD2 of DISP1 (Fig. 4B), and includes hydrophobic contacts and several possible ionic interactions with total buried surface area of 989.3 Å^2^ (−8.6 kcal/mole, calculated using PISA[30]). An additional striking feature of the interaction involves the furin-cleaved front-side linker of DISP1-A, which extends from ECD2 upward and around ShhN in a one-armed molecular embrace (Fig. 4A). The DISP1-A density corresponding to this linker was considerably sharper than in the DISP1-A apostructure, allowing us to extend the model of the linker by an additional 13 residues preceding the furin cleavage site (Fig. S2). The region of ShhN contacted by this linker arm corresponds to a polybasic region previously shown to interact with heparin sulfate or other sulfated glycosaminoglycans (GAGs)[31], which would be displaced by this molecular embrace. A sharpening of the DISP1-A density within the ShhN:DISP1-A map also allowed us to extend the DISP1-A model by an additional 11 residues (399-409) in the distal portion of ECD1 (Fig. 4B; Fig. S2).

### Conformational dynamics of DISP1

We noted a somewhat more compact conformation of ECD1 and ECD2 of DISP1-A in its pincer-like grasp of ShhN, similar in many respects to the T conformation identified as the second major subclass in 3D classification of the apoprotein preparation (Fig. S5). To more systematically investigate the range of DISP1-A conformations, we applied 3D variability analysis (3DVA) [32] to the dataset collected with the addition of ShhN. The first principal component (PC1) of this analysis revealed a range of conformations anchored at one end of the reaction coordinate by a ShhN:DISP1-A complex like the one just described, and the opposite end by a DISP1-A molecule lacking the ShhN ligand, and with the furin-cleaved linker arm no longer resolved (Fig. 5A, Supplementary Video S1). We similarly applied 3DVA to the dataset from which our initial DISP1-A structure was derived, and found within PC1 of this dataset a similar range of DISP1-A conformations, with ECD changes similar to those seen in PC1 of the complex preparation (Fig. 5). We interpret the similarity of conformations along this reaction coordinate, with ShhN binding restricted to one end, as resulting from selective binding to a restricted subset of possible DISP1-A conformations.

**Fig. 5.**
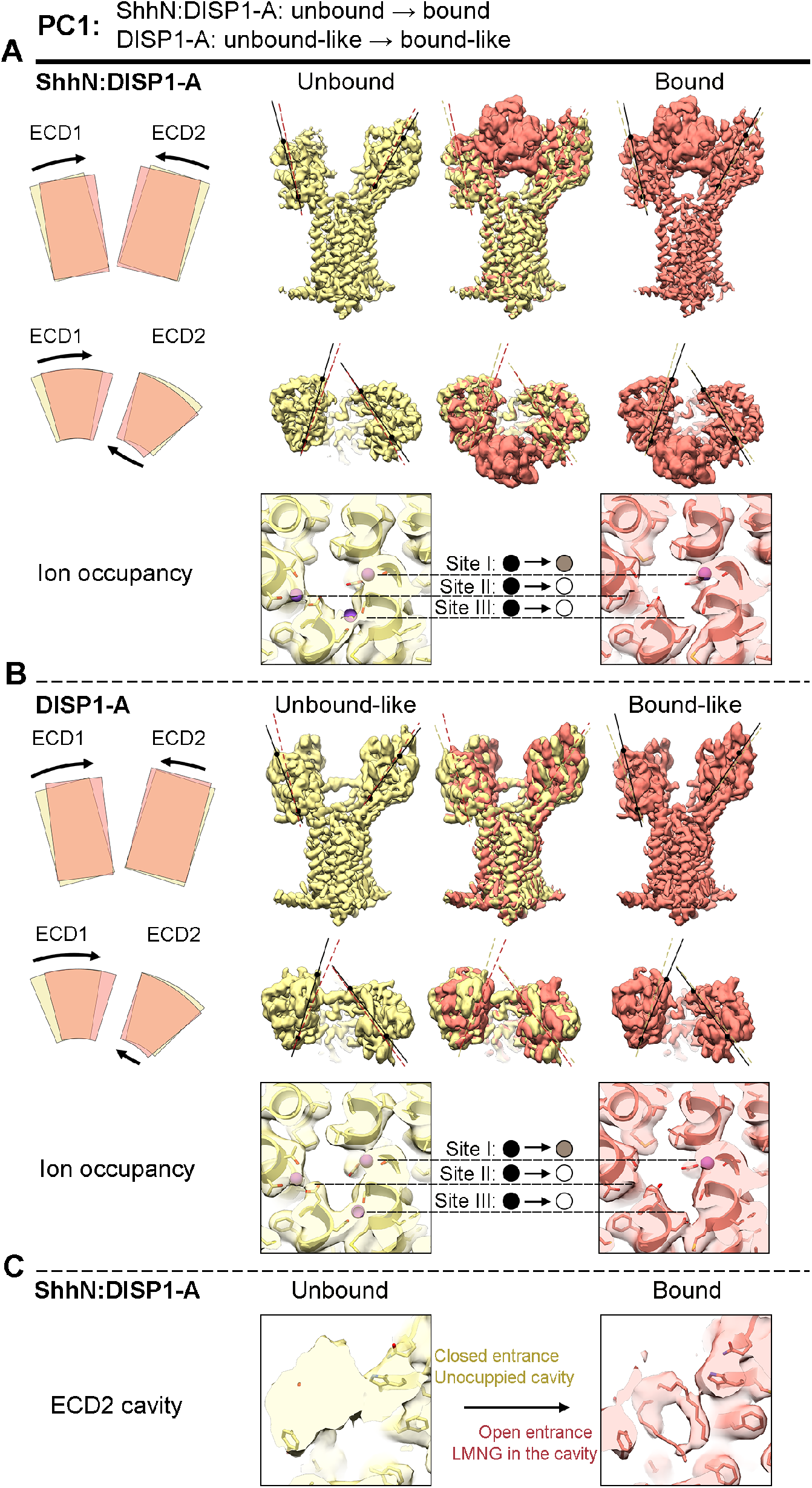
Conformational dynamics link intramembrane Na^+^ site occupancy to Hedgehog release. **A**, Three-dimensional variability analysis (3DVA) of the dataset with added ShhN revealed a series of DISP1-A conformations along the first principal component (PC1) revealed ShhN bound or absent at opposite ends. Reconstructed densities from the extremes of PC1 (unbound, khaki; bound, salmon) are shown from the side and the top with a superimposed view of these extremes in the center. The changes in ECD position are illustrated by lines drawn atop the reconstructed densities and by the schematized shapes and arrows to the left. Reconstructed densities in the ion coordination nexus in the transmembrane domain indicate that Na^+^ site occupancy also changes, from fully occupied in the unbound state to site I only occupied in the bound state. See also Supplementary Video S1. **B**, Similar views of the apoprotein prep show a very similar set of changes in reconstructed densities from PC1 of 3DVA, along with similar shifts in the occupancy of the Na^+^ ion sites. These similarities suggest that a tight linkage between Na^+^ site occupancy and DISP1 conformation determines binding and release of the ShhN signal. **C**, From the 3DVA in **A**, the hydrophobic cavity at the distal end of ECD2 is occupied by an LMNG-like density in the bound state, but is unoccupied and has its entrance blocked by ingress of F846 and other nearby residues in the unbound state. See Supplementary Video S2.

Interestingly, a close examination of the transmembrane domains within the dataset with added ShhN reveals that at a given contour level, the Na^+^ sites are differentially occupied at opposite extremes of the PC1 reaction coordinate (Fig. 5A, Supplementary Video S1). We thus see densities corresponding to all three Na^+^ ions at the end of the reaction coordinate representing the unbound DISP1-A conformation, whereas only Na^+^ site I appears to be occupied at the opposite end of the reaction coordinate, representing the ShhN-bound conformation. A similar differential ion site occupancy is also noted at extreme ends of the PC1 reaction coordinate for the DISP1-A apoprotein preparation (Fig. 5B), suggesting that Na^+^ ion occupancy drives the conformational state of DISP1 whether ShhN is present or not; changes in Na^+^ ion occupancy occasioned by chemiosmotically driven flux via the DISP1 transmembrane tunnel thus could drive a dynamic cycle of conformational changes that cause Hedgehog binding and release.

### Extraction and release of Hedgehog lipid adducts

Because release of the mature Hedgehog signal would require extraction of its covalently linked cholesteryl and palmitoyl adducts from the membrane, several features of our structures are worth noting. First, the lifted sterol on the front side of the DISP1-A (Fig. 1E,F) may correspond to an intermediate state of the cholesteryl adduct as it is removed from the hydrophobic environment of the membrane during ShhNp release. Second, the hydrophobic cavity at the distal tip of ECD2 (Fig. 1E,F) is large enough to easily accommodate one of the covalently linked lipid adducts. Interestingly, in our 3DVA of the preparation with added ShhN, we note that this cavity is closed off in the ShhN-unbound state by ingression of the side chain of F846, and the non-proteogenic density is absent (Fig. 5C, Supplementary Video S2).

The possible involvement of the ECD2 distal hydrophobic cavity in final release of modified Hedgehog raises the question of how a Hh-linked lipid might reach the distal ECD2 cavity in the first place. The PTCH1 protein employs an enclosed hydrophobic conduit formed by the juxtaposition of ECD1 and ECD2 to execute a similar movement of sterols away from the membrane. In DISP1, although ECD1 and ECD2 are split apart in a manner that would bisect this conduit, the ECD2 remnant retains a series of hydrophobic residues that line its inner surface, at the edge of the ferredoxin-like α+β sandwich fold. This surface could perhaps form a partial hydrophobic conduit from the membrane to the hydrophobic cavity near the distal tip of ECD2 (Fig. 1F), analogous to the sterol conduit within the center of the conjoined ECDs of PTCH1.

## Discussion

How does DISP1-A extract and release the lipid-modified Hedgehog signal from the membrane? We propose that this process requires an activated state of ShhNp, features of which are represented by our ShhN:DISP1-A complex structure, followed by release (Fig. 6). We suggest that during activation the favorable interaction energy of DISP1-A binding to ShhN, available only at a location ~25 Å above the membrane, helps offset the unfavorable energy of membrane extraction of the Hh-linked lipids. In addition, the lifted sterol at the front side of DISP1-A may represent an intermediate in cholesteryl adduct extraction, and the hydrophobic streak leading to the distal hydrophobic cavity in ECD2 (Fig. 1F) may represent a path to an energetically neutral way-station for one of the lipid adducts. This activated form of DISP1 bound to the Hedgehog protein is characterized by three prominent features (Supplementary Videos S1-2): (*i*) a pincer-like action of the split ECDs; (*ii*) an embrace of ShhN mediated by the furin-cleaved linker; and (*iii*) accommodation of a lipid adduct in distal ECD2. From this activated state, which is tightly correlated with restriction of Na^+^ occupancy to site I, the availability of SCUBE2 and the force of Na^+^ flux down its chemiosmotic gradient together drive release of ShhNp, accompanied by opening of the ECD pincers, expulsion of the ECD2 lipid, withdrawal of the linker embrace, and resetting to permit another cycle. The position of ShhN relative to DISP1-A in our complex structure is distinct in its location, orientation, and interaction interface from the position recently proposed by Cannac et al.[33] for *Drosophila* HhN, based on docking of a *Drosophila* HhN structure[34] within a 4.8 Å density map (Fig. S9). This difference is somewhat puzzling in light of the ability of mammalian DISP1 to rescue *Drosophila disp* mutant function^6^, and we cannot definitively account for it. One functional difference is that mammalian DISP1 cooperates with SCUBE2 for its Hedgehog-releasing activity, whereas *Drosophila* lacks a Scube orthologue.

**Fig. 6.**
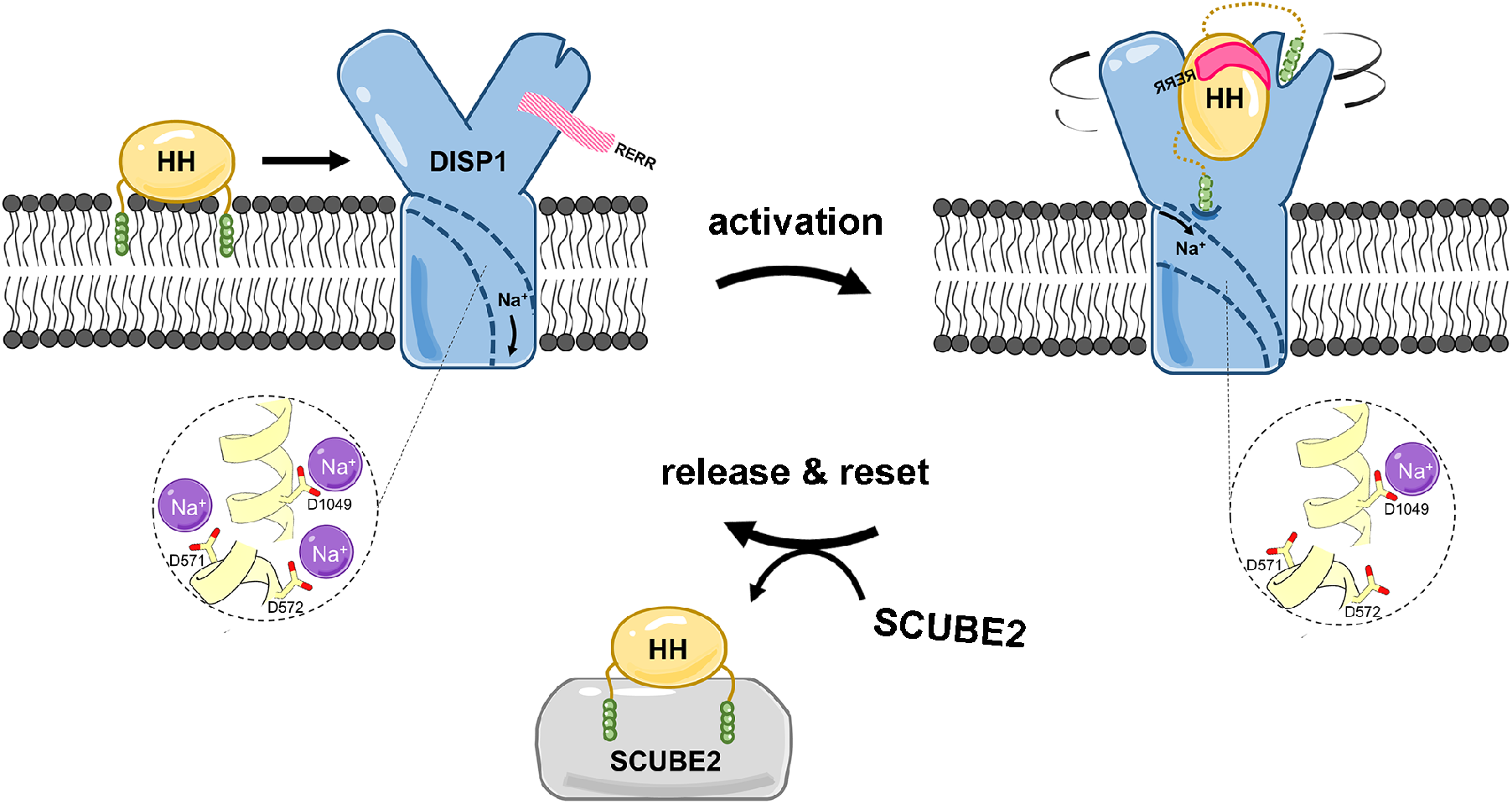
A model for Na^+^-driven conformational dynamics in the DISP1 activity cycle. Membrane anchored, dually lipid-modified Hedgehog signal (left side), is activated for release by interaction with DISP1 ECDs, at a location ~25 Å above the membrane. Hedgehog binding and activation (right side) entails (*i*) a pincer-like action of the DISP1 ECDs; (*ii*) interaction with and stabilization of the DISP1 furin-cleaved linker (red); and (*iii*) accommodation of a Hedgehog-linked lipid in the distal cavity of DISP1 ECD2. From this activated state, associated with restricted Na^+^ occupancy (site I only), SCUBE2 availability and the chemiosmotic force of the transmembrane Na^+^ gradient together drive hedgehog release and resettng of DISP1 for resumption of the cycle (left). See Supplementary Videos S1 and S2.

We note from structure-based alignment that the position of each of 15 protein-derived Na^+^-liganding oxygens in DISP1 (10 from side-chains, five from main-chain carbonyls) is conserved in PTCH1, as are the three charge-neutralizing intramembrane acidic residues (Fig. S10). This degree of conservation strikingly exceeds that of the overall protein sequence (19% identity overall), and strongly supports the functional role of Na^+^ flux in DISP1, as PTCH1 has been shown to suppress SMO activity in a Na^+^ gradient-dependent manner[25]. All of these residues are also conserved by sequence alignment in PTCH2, which also functions to suppress SMO[35]. In contrast, DISP2, which unlike DISP1 fails to rescue *disp* function in *Drosophila[10]* and probably lacks Hedgehog-releasing activity, has only 1 of three intramembrane acidic residues and 6 of 10 side-chain oxygens. Other mammalian family members, PTCHD2 and PTCHD3, display complete conservation of charge-neutralizing and liganding oxygen side-chains and thus seem likely to utilize a Na^+^ gradient in their functions, whereas PTCHD1 and PTCHD4 show poor conservation of these residues and might have dispensed with transport activity or adopted a different mechanism for sustaining it. An intermediate level of conservation is seen for NPC1, which functions in the lysosome and thus has available to it both Na^+^ and H^+^ gradients[36]. Prokaryotic RND transporters harbor distinct residues at these Na^+^-coordinating sites, consistent with their predominant utilization of proton motive force across the bacterial inner membrane to power their transport activities (Fig. S10). Despite this lack of conservation, some prokaryotic transporters, such as the SecDF1 protein in halophiles (Fig. S10), nevertheless require Na^+^ for their activities, suggesting that coupling of Na^+^ flux to transport may have evolved more than once.

How do DISP1 and PTCH1, which share transmembrane topology, conserved sequence elements, and hence a common evolutionary origin, and which both interact with lipid-modified Hedgehog protein, produce such strikingly distinct functional effects on Hedgehog pathway activation? Our analysis suggests an underlying mechanistic similarity, namely, the use of chemiosmotic force from the metazoan transmembrane Na^+^ gradient to power the release of lipids from membranes. In the case of PTCH1, cholesterol removal *via* an enclosed sterol conduit in the PTCH1 ECD regulates SMO by depriving it of the sterol that must be bound within the interior of its 7 transmembrane helical bundle for activation[37,38]. The central activity of DISP1 also appears to be the extraction of lipids from the membrane, but in this case the lipids are covalent adducts of the Hedgehog protein signal, thus releasing Hedgehog for its patterning activity and necessitating a splitting of the DISP1 ECD to accommodate the protein moiety. This ECD domain splitting and the unusual interaction of the furin-cleaved linker arm with Hedgehog at a position elevated above the membrane together help explain the requirement for furin cleavage in DISP1 function[23].

Remarkably, acquisition of both the furin cleavage site and the preceding linker arm that contacts ShhNp appears to have been facilitated in evolution by the insertion into the first extracellular loop of DISP1 of a single small exon that encodes both features (Fig. S2). Whereas this exon insertion contributes to functional divergence between DISP1 and PTCH1, a fundamental mechanistic similarity is indicated by a lifted sterol in PTCH1, at a position within the 12TM bundle near the beginning of the sterol conduit[39]. The unusual positions of these sterols at corresponding locations in both proteins suggest partial extraction from the membrane, perhaps representing a common intermediate in the lipid extraction activities of both proteins.

## Materials and Methods

### DISP1 expression and purification

All constructs were cloned using Gibson Assembly. In DISP1-A, DISP1-A-NNN, and the ion-site mutants in Fig. 2, amino acids 3 to 171 were deleted from the full-length (1-1521) DISP1. For protein expression, DISP1-A:ShhNp binding assays and ShhNp release assays, different DISP1 variants were cloned into the pcDNAh vector[17].

For detection of protein expression levels in Fig. S1D, HEK293T cells were seeded into a 6-well plate. Plasmids encoding DISP1 and DISP1-A with an HA tag at the C-terminus were transiently transfected into the cells using Lipofectamine 3000 (ThermoFisher). Two days after transfection, the cells were lysed in buffered detergent. Protein extracts were cleared by a subsequent centrifugation (100,000 xg, 30 min, 4°C). Western blotting was performed using Criterion TGX stain-Free precast gels (Bio-rad) and Immobilon-P PVDF membrane (Millipore). PVDF membranes were immunoblotted overnight at 4°C with anti-HA rabbit polyclonal (1:1000, Cell signaling) or anti-β-actin rabbit polyclonal (1:1000, Cell signaling) antibodies followed by a 1-hour incubation with the corresponding HRP-conjugated secondary antibodies (1:2000, Promega) at room temperature. The blots were developed using SuperSignal West Pico PLUS substrate (ThermoFisher).

For large-scale protein purification, C-terminally SBP-tagged DISP1-A was cloned into a BacMam expression vector, PVLAD6 vector. Baculoviruses were produced as previously described[17,25]. HEK293F cells were cultured in suspension using Free-Style medium (ThermoFisher) until the density reached 1.5 – 2 ×10^6^ cells/ml, and cells were transduced with DISP1-A-SBP baculoviruses in the presence of 10 mM sodium butyrate (Sigma). Forty-eight hours after transfection, cells were collected and stored at −80°C. For DISP1-A-NNN, plasmids encoding DISP1-A-NNN-SBP were transfected into the HEK293F cells using linear polyethylenimines (PEIs, Polysciences). Both DISP1-A and DISP1-A-NNN were purified essentially as previously described[17]. The C-terminally SBP-tagged DISP1-A or DISP1-A-NNN was captured using Streptavidin-agarose affinity resin (ThermoFisher). The eluted protein was concentrated by an Amicon Ultra filter (100 kDa cutoff, Millipore) and further purified using Superose 6 Increase 10/300 GL column (GE Healthcare).

### ShhN expression and purification

DNA fragment encoding mouse Sonic hedgehog residues 26-189 was cloned into a pHTSHP vector with N-terminal His_8_ tags.[29] Protein was expressed in *E*. coli strain BL21(DE3) and purified as previously described in ref. [17]. The cell lysate was first centrifuged at 8,000 xg for 15 min (4°C), then the filtered supernatant was applied to the Ni-NTA resin. After elution from the Ni-NTA column, the N-terminal tags were cleaved with HRV3C protease. The protein was dialyzed against 20mM HEPES, 100mM NaCl, and 7mM β-mercaptoethanol, and further purified by SP Sepharose cation exchange chromatography.

### Cryo-EM sample preparation and data acquisition

For DISP1-A, 2.5 μl of purified and concentrated sample at a concentration of 1.56 mg/ml was applied to glow-discharged gold grids covered with holey carbon film (Quantifoil, 300 mesh 1.2/1.3) and plunge frozen using a Vitrobot Mark IV, with blotting time 4 seconds at 8°C and 100% humidity. DISP1-A-NNN cryo-EM grids were prepared similarly, but using gold-grids covered by a holey gold film (Quantifoil UltrAufoil, 300 mesh 1.2/1.3) and a sample concentration of 0.3 mg/ml. For ShhN:DISP1-A complex, purified DISP1-A was incubated with ShhN for 3 hours (4°C) before cryo-sample preparation. DISP1-A:ShhN grids were prepared in the same way as that for DISP1-A.

Cryo-EM data for DISP1-A and ShhN:DISP1-A were collected with a Titan Krios microscope operated at 300keV and equipped with a Bio Quantum post-column energy filter with zero-loss energy selection slit set to 20eV. Using SerialEM [40], images were recorded automatically at a nominal magnification of 105 (true magnification 59,880X), with defocus range set to −0.5 to −1.5 μm. Images were recorded with a post-GIF Gatan K3 camera in super-resolution single electron counting mode with a calibrated pixel size of 0.4175 Å (physical pixel size of 0.835 Å) The DISP1-A-NNN samples were imaged on a different Titan Krios with identical parameters, but a slightly different calibrated pixel size of 0.417 Å. Data collection statistics are shown in Table S1.

### Imaging processing and map calculation

As part of an on-the-fly image processing pipeline, which processes micrographs immediately after acquisition, we used MotionCor2[41] to correct motion of each dose fractionated movie stack, sum motion corrected frames with and without dose weighting, and bin images in Fourier space to the physical pixel size (~0.835 Å) for further image processing. Motion-corrected, dose-weighted sums were used for contrast transfer function determination and resolution estimation in cryoSPARC[27]. Particles were picked in cryoSPARC first using a Gaussian template, and then re-picked with low-resolution templates generated from an initial structure. Subsequent image processing was carried out in RELION[42], cryoSPARC and cisTEM[43]. The flow-chart of image processing is illustrated in Fig. S3 (DISP1-A and DISP1-A-NNN) and Fig. S8 (ShhN:DISP1-A). Conversion of data from cryoSPARC to Relion and cisTEM was performed using UCSF pyem[44]. Directional Fourier shell correlation curves were determined as previously described[45].

### Model building and refinement

The *de novo* atomic models of DISP1-A were built in Coot[46], refined in PHENIX[47] and finalized in ISOLDE[48]. Starting from a predicted model from PHYRE2[49] and a combined density of R and T conformers, we manually fitted transmembrane (TM) helices into the corresponding density and then traced the backbone of ECDs and linkers in between those TM helices in Coot. The preliminary model was then mutated based on Mouse DISP1 sequence and refined using real space refinement in PHENIX. Based on residual unmodeled density near the protein, ligands were added and refined in real space using Coot and PHENIX, respectively. Starting from this “common model” built referring to the combined density of R and T, we obtained models for each conformer (R, T and NNN) by refining this “common model” against the final density of each conformer individually, removing atoms unresolved in each density and then finalizing the models with ISOLDE (largely to improve steric clashes). The ShhN:DISP1-A complex was treated in the same fashion, but using several crystal structures of mammalian Shh as initial models (PDB codes: 1vhh, 4c4n, and 6pjv). All models were validated by wwPDB validation server[50] and no major issues were reported in our three models.

Visualizations of the atomic models (figures) were made using UCSF Chimera[51], ChimeraX[52] and PyMOL (The PyMOL Molecular Graphics System, Version 2.0 Schrödinger, LLC.). The ion conduction path was mapped using Caver 3[26] and ion coordination geometries analyzed with the *CheckMyMetal* webserver[24]. The ESPript server[53] was used in preparation of the sequence alignment.

### DISP1-ShhNp binding assay

The DISP1-ShhNp binding assay was conducted using the HEK293 Flp-In T-REX ShhNp-producing cell line[15]. ShhNp-producing cells were maintained in DMEM (ThermoFisher) with 10 % FBS (Omega) and 1% Penicillin-Streptomycin-Glutamine (ThermoFisher). Plasmids of C-terminally HA-tagged DISP1-A variants were transiently transfected into the cells using Lipofectamine 3000. One day after transfection, full-length Shh expression was induced by 4 μg/ml Doxycycline (Sigma) in DMEM F-12 (ThermoFisher) supplemented with 1% Insulin-Transferrin-Selenium (ThermoFisher) and 1% Penicillin-Streptomycin-Glutamine. Another day after ShhNp expression was induced, cells were lysed in buffered detergent. Following a subsequent centrifugation (100,000 xg, 30 min, 4°C), Streptavidin-agarose affinity resin was used to pull down the SBP-tagged DISP1-A variants, together with any bound ShhNp, from the supernatant. Immunoprecipitates were analyzed by Western blot as described above using the following antibodies: anti-HA rabbit polyclonal (1:1000) and anti-Shh rabbit monoclonal (1:1000, Cell signaling).

### ShhNp release assay

Disp^−/−^ Mouse Embryonic Fibroblasts (MEFs)[10,15], were cultured on 6-well plates in DMEM supplemented with 10 % FBS and 1% Penicillin-Streptomycin-Glutamine. All the cell transfections were performed using Lipofectamine 3000. Plasmids encoding C-terminally GFP-tagged (Fig. S1E) or HA-tagged (Fig. 2E) DISP1 variants were transfected into the cells along with full-length Shh constructs. The ratio of plasmids expressing DISP1 variants and Shh was 1 to 2. For conditions that did not have DISP1 variants or Shh, GFP constructs were used as a negative control to replace the missing partner. Two days after transfection, following a quick wash with DMEM, the cells were shifted to SCUBE2 conditioned medium or control medium (600 μl/well). After 16 hours of incubation, the medium was collected and centrifuged at 18,000 xg for 30 min (4°C). The supernatant was probed by immunoblotting for ShhNp (anti-Shh rabbit monoclonal, 1:1000), 3xFlag-tagged SCUBE2 (anti-Flag m2 mouse monoclonal, 1:1000, sigma). Cells were lysed using the same buffer as that in DISP1-ShhNp binding assays (400 μl/well). Cell lysate was probed by immunoblotting for ShhNp, HA or GFP-tagged DISP1 variants (anti-HA rabbit polyclonal, 1:1000 or anti-GFP rabbit polyclonal, 1:2000, Invitrogen), and β-actin (anti-β-actin rabbit polyclonal, 1:1000).

SCUBE2 conditioned medium was prepared using the HEK293 Flp-In T-REX Scube2-3xFlag-producing cell line, as previously described[15]. Briefly, SCUBE2-producing cells were grown in DMEM with 10 % FBS and 1% Penicillin-Streptomycin-Glutamine. Four μg/ml Doxycycline (Sigma) was used to induce SCUBE2-3xFlag expression in DMEM F-12 (ThermoFisher) supplemented with 1% Insulin-Transferrin-Selenium (ThermoFisher) and 1% Penicillin-Streptomycin-Glutamine. Two days after induction, Scube2 conditioned medium was collected, cleaned by centrifugation (1,000 xg, 10 min, 4°C) and filtered by a 0.22 μm filter. The control medium was prepared using a medium formulation identical to that for SCUBE2 conditioned medium.

## End Matter

### Author Contributions and Notes

Q.W. purified and characterized DISP1-A, DISP1-A-NNN and ShhN:DISP1-A complex and performed ShhNp binding and release assays. D.E.A. prepared cryo-EM grids and collected and processed cryo-EM data. D.E.A. also performed the 3D variability analysis. K.D., D.E.A., Y.Z., and Q.W. built the models. Y.M. provided mouse genetic epistasis analysis while in the laboratory of P.A.B. at Johns Hopkins University. All authors participated in discussion and analysis of the data. P.A.B., Q.W., D.E.A., K.D., Y.C., and Y.Z. prepared the manuscript.

The authors declare no competing interests.

This article contains supporting information: Supplementary Figures S1–10

Supplementary Table S1

Supplementary Videos S1 DISP1 protein conformation and Hedgehog binding/release are linked to Na^+^ site occupancy.

Animations of the first principal component identified by 3D variability analysis of ShhN:DISP1-A cryo-EM data, containing bound and unbound DISP1-A particles. In three synchronous views, the left shows a wide-angle view from the front of DISP1 (khaki), with bound ShhN (goldenrod) appearing in coordination with tensing of DISP1 ECDs. The center view highlights at a lower density threshold the DISP1 residues preceding the furin cleavage site, which form a one-armed embrace of ShhN upon binding (hot pink). Right, a tilted, cut-away top view illustrating loss of density in transmembrane Na^+^ coordination sites II & III, again coincident with ShhN binding.

Supplementary Video S2. Conformational dynamics of the ECD2 hydrophobic cavity.

Left, animation of the first principal component from 3DVA of ShhN:DISP1-A, with perspective from the back of DISP1-A, looking over ECD1 towards ECD2, and with the region of the hydrophobic pocket in ECD2 highlighted in blue. In the bound state, the pocket is open, and filled by a clear lipidic density (likely LMNG), while in the unbound state the pocket is closed by ingress of F846 and other nearby residues. Right, synchronized cut-away view of the Na^+^ coordination sites, reproduced from Supplementary Video S1.

## Acknowledgments

We thank K. Roberts, R. K. Mann, and A. Kershner for technical assistance and advice, and V. Korkhov and X. Li for sharing data and information prior to publication. Y.C. is an investigator of the Howard Hughes Medical Institute. Supported by NIH grants R01GM102498 (to P.A.B.) and R01GM098672, S10OD020054, P50AI15476 and S10OD021741 (to Y.C.).

---

**Fig. S1.**
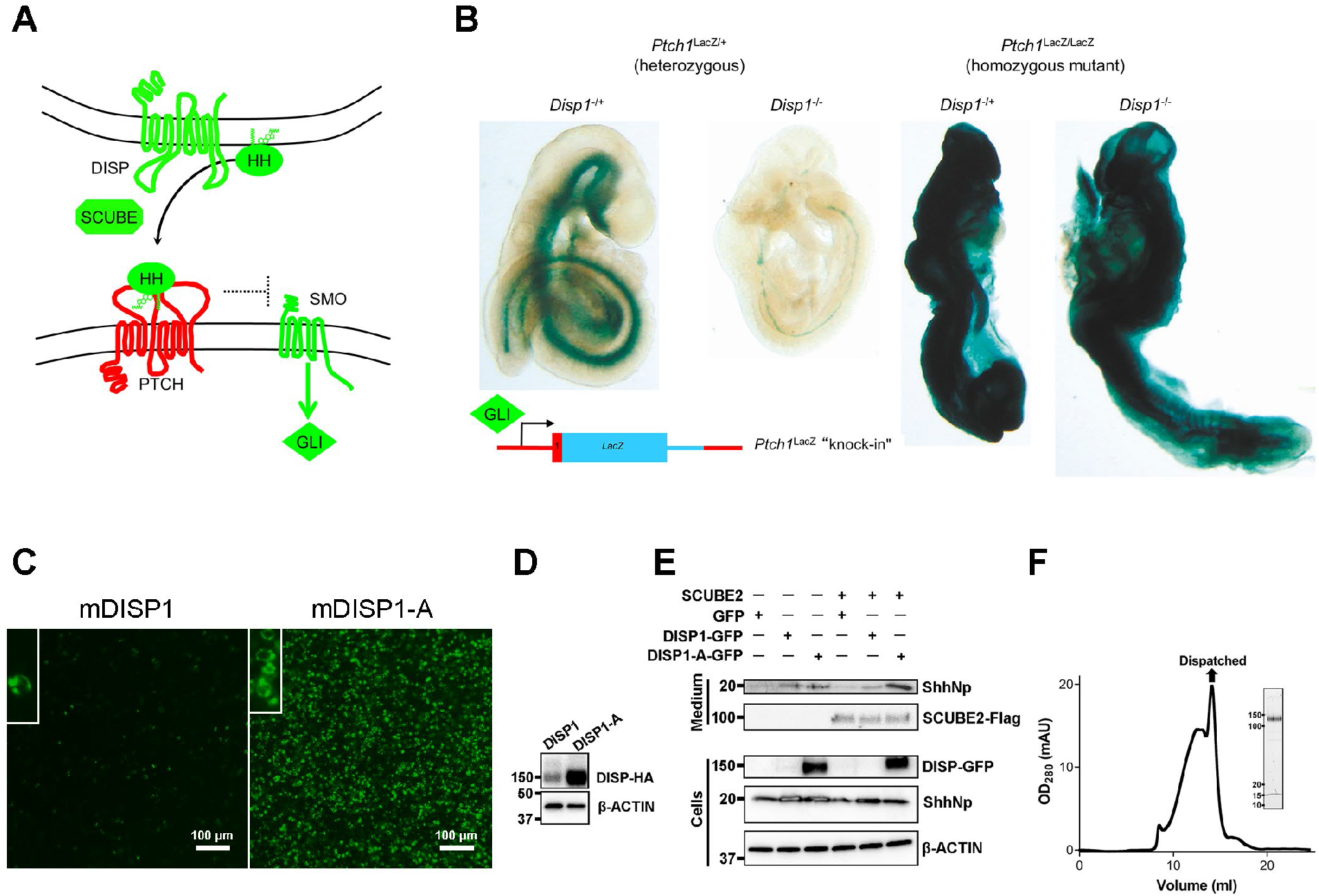
Dispatched function and expression. **A-B**, Opposing functions of structurally related transporters Dispatched and Patched in Hedgehog signaling. **A**, The HH protein signal, covalently modified by cholesterol and palmitate, requires the action of DISP1 and SCUBE for release from the membrane of producing cells. HH then uses its palmitoyl adduct to clog the sterol transport conduit and block the function of its receptor PTCH1 in responding cells. The loss of PTCH1 sterol transport activity permits accumulation of cholesterol within the inner leaflet to levels that activate SMO by binding within its seven transmembrane helical bundle, resulting in activation of the GLI transcriptional effector of Hedgehog signaling. **B**, As *Ptch1* is a target for GLI activation, the X-Gal staining of a *Ptch1^LacZ^* knock-in allele[22] provides an indication of Hedgehog pathway activity (leftmost embryo). Homozygous mutation of *Disp1*[10] causes a loss of nearly all embryonic Hedgehog pathway activity (2^nd^ embryo from left), whereas homozygous disruption of *Ptch1* leads to unregulated ectopic pathway activity, regardless of the functional status of *Disp1* (rightmost two embryos). **C**, DISP1-GFP and DISP1-A-GFP in HEK293T cells with magnified insets showing expression on the cell membrane. **D**, Western blot showing high level of DISP1-A expression relative to DISP1 (both proteins HA-tagged). **E**, Functional assay in *Disp^−/−^* mouse embryonic fibroblasts (MEFs). Culture media and cell lysates from transiently transfected *Disp^−/−^* MEFs were probed by immunoblotting for expression of DISP-GFP, ShhNp, and SCUBE2. DISP1 and DISP1-A both released ShhNp in the presence of SCUBE2. Panels **D** and **E**, show representative results (n = 4); β-ACTIN, loading control. **F**, Size exclusion chromatography of purified DISP1-A, together with the SDS-PAGE of the indicated fraction, corresponding to monomeric DISP1-A.

**Fig. S2.**
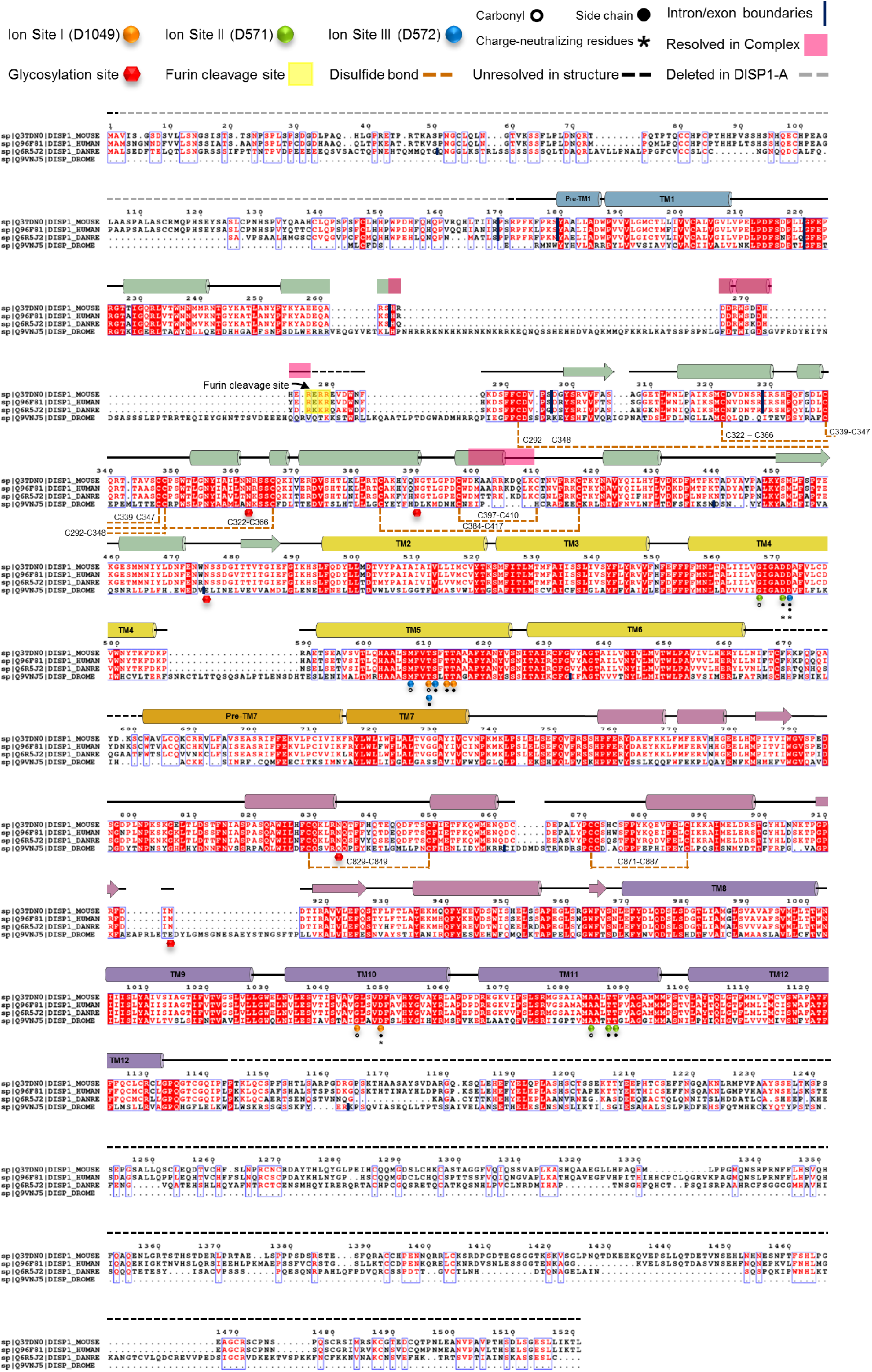
Sequence alignment. Sequence alignment of Dispatched proteins from *Mus musculus* (mouse), *Homo sapiens* (human), *Danio rerio* (zebrafish) and *Drosophila melanogaster* (fruit fly). Amino acid sequences of the structurally determined DISP1-A protein are indicated. Secondary structure elements are indicated above the sequence, with subdomains colored according to Fig. 1A-B, and other features indicated as shown above. The ESPript server[53] was used in preparation of the sequence alignment. (TM, transmembrane helix.)

**Fig. S3.**
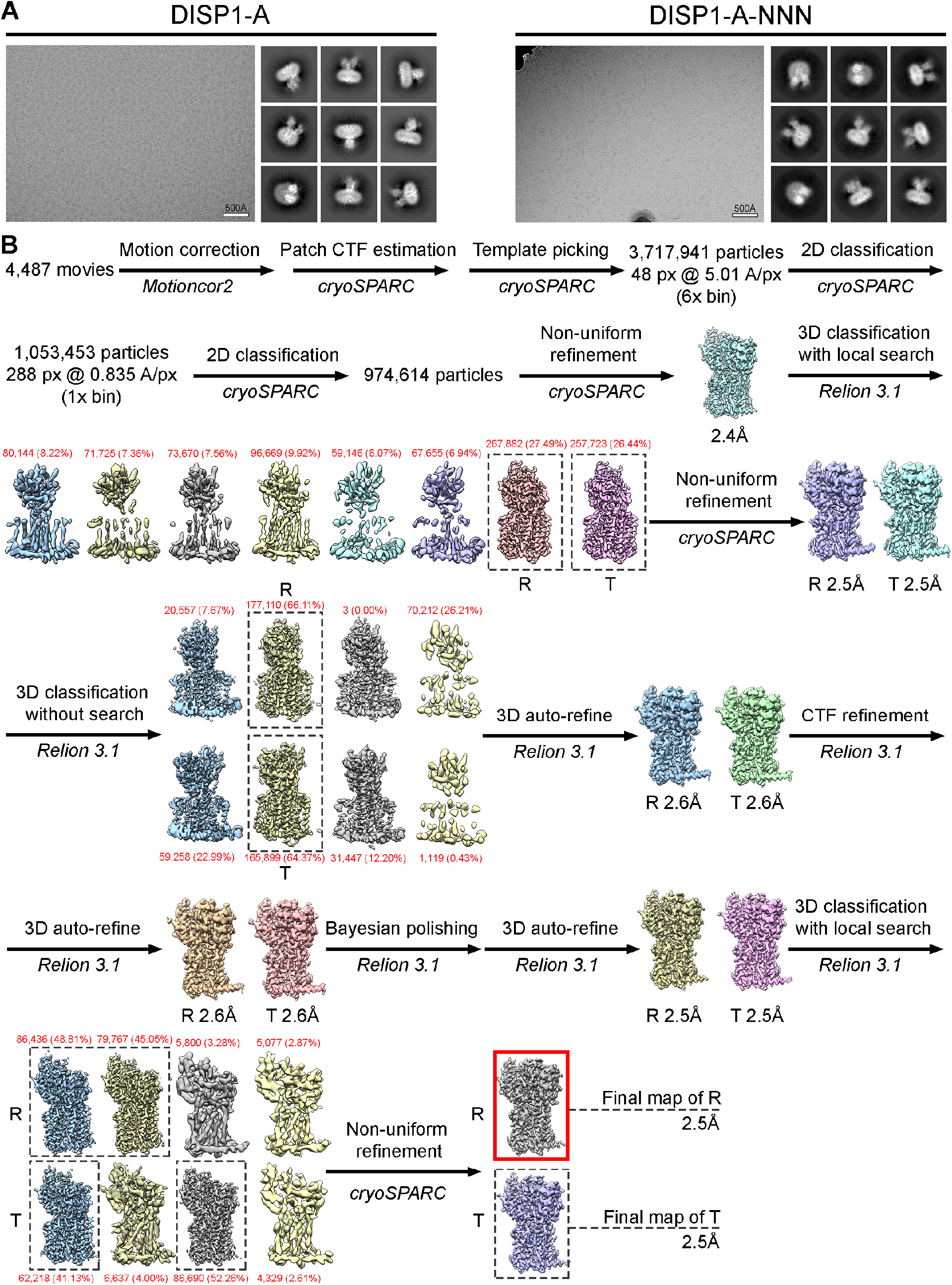
Cryo-EM data and image processing flow for DISP1-A and DISP1-A-NNN. **A**, A representative cryo-EM micrograph and several highly-populated, reference-free 2D class averages are shown for DISP1-A (left) and DISP1-A-NNN (right). The micrograph for DISP1-A-NNN has been contrast-stretched for display in order to account for the presence of a gold edge in the upper left corner of the image (the DISP1-A-NNN particle distribution on this grid necessitated targeting of the hold edge). **B**, Schematic flow-chart representing the image processing approach for DISP1-A. Thumbnail images of each 3D class or refinement are shown along with global GS-FSC resolution in black, particle counts in red, and dashed black boxes to indicate selected 3D classes. After separation of the R and T conformations in the first round of 3D classification, the identical processing flows for the two conformations are shown in parallel. Cryo-EM map (Red box) and atomic model of R conformation are used in main figures to present DISP1-A features.

**Fig. S4.**
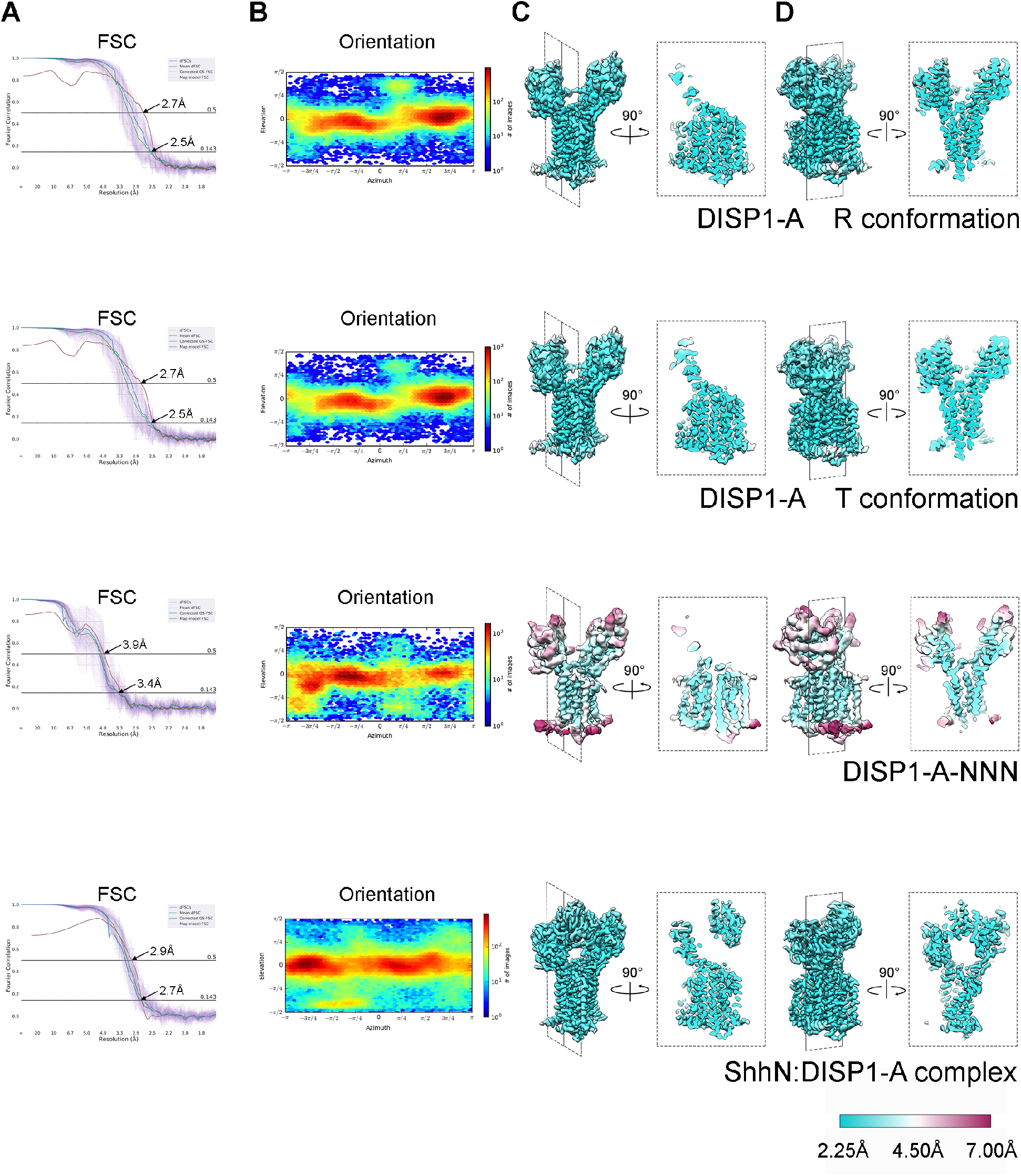
Cryo-EM density and atomic model quality. Fourier shell correlation curves (**A**), particle orientation distributions (**B**), and local resolution maps (**C,D**) are shown for R and T conformations of DISP1-A, DISP1-A-NNN, and for ShhN:DISP1-A complex. The “gold-standard” independent half-map FSC curves and orientation distributions were determined during refinement in cryoSPARC, map-to-model FSC curves were calculated n PHENIX using protein chains only, and directional FSC curves were estimated as in ref. [45]. The orientation distributions are plotted such that an elevation angle of 0° corresponds to a “side-view” perpendicular to the transmembrane helices; in each case the predominant views are “side-views” at a wide range of azimuthal angles. Local resolution estimates were computed using the BLOCRES algorithm as implemented in cryoSPARC.

**Fig. S5.**
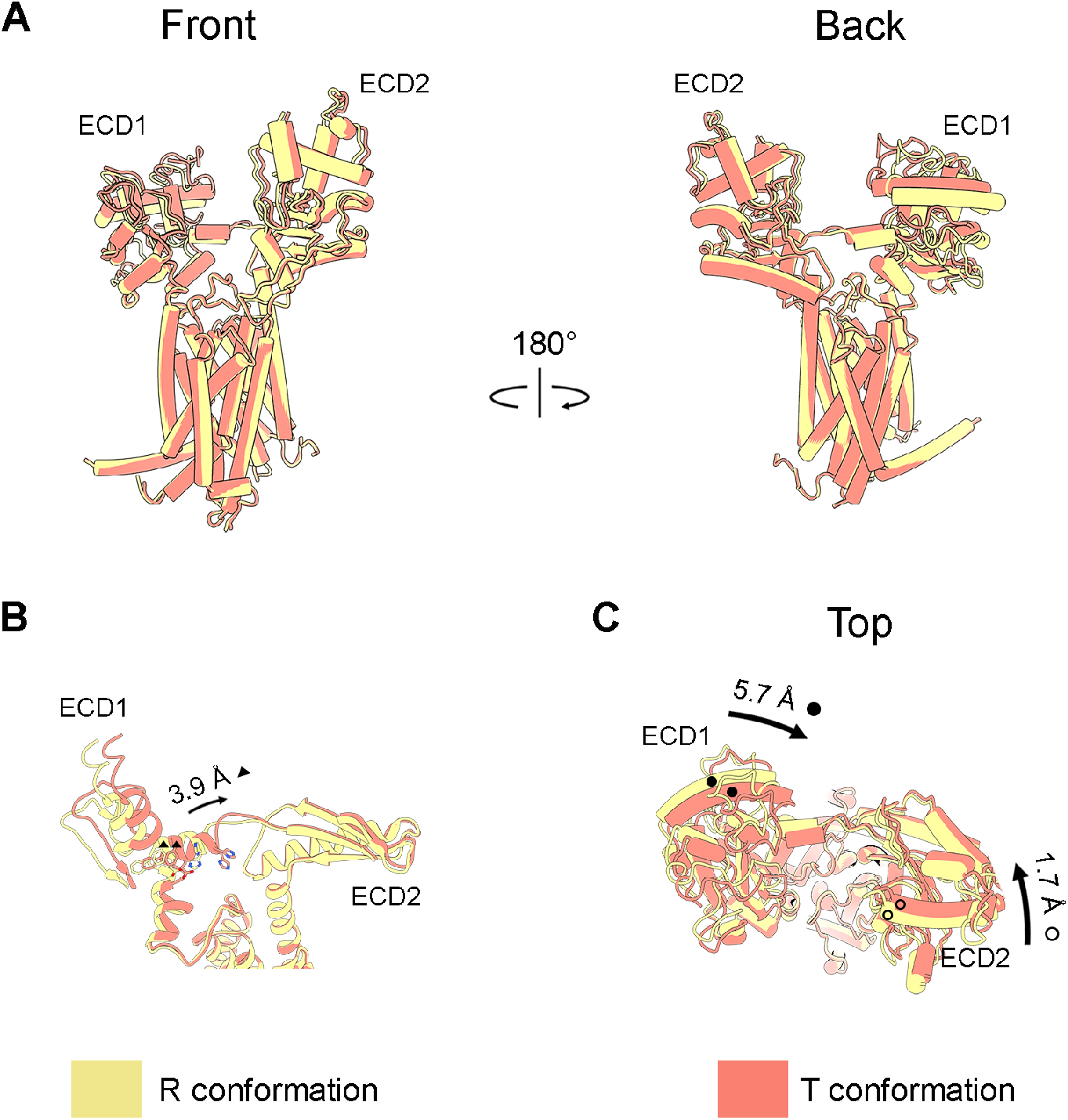
DISP1 conformational flexibility revealed by 3D classification. **A**, Overlay comparing R and T conformations (khaki and salmon, respectively), including front and back views. Major conformational changes are localized to the extracellular domains. **B**, Cut-away view showing the formation of a “kink” in the back-side linker of the T conformation, with an accompanying shift of about one helix turn that breaks a hydrogen bond between linker residue H777 and the backbone carbonyl of K767. **C**, View of the R and T conformations from the extracellular side, highlighting the movement of secondary structure elements in ECD1 (> 5 Å) and ECD2 (~2 Å). The shift of ECD1 and the formation of the inter-ECD linker “kink” appear intimately related. Numbers indicate distances (Å) between the Cα of F772 in R and T (marked by ▲ in **B**) and R382 and S898 (marked by ● and ○, respectively, in **C**).

**Fig. S6.**
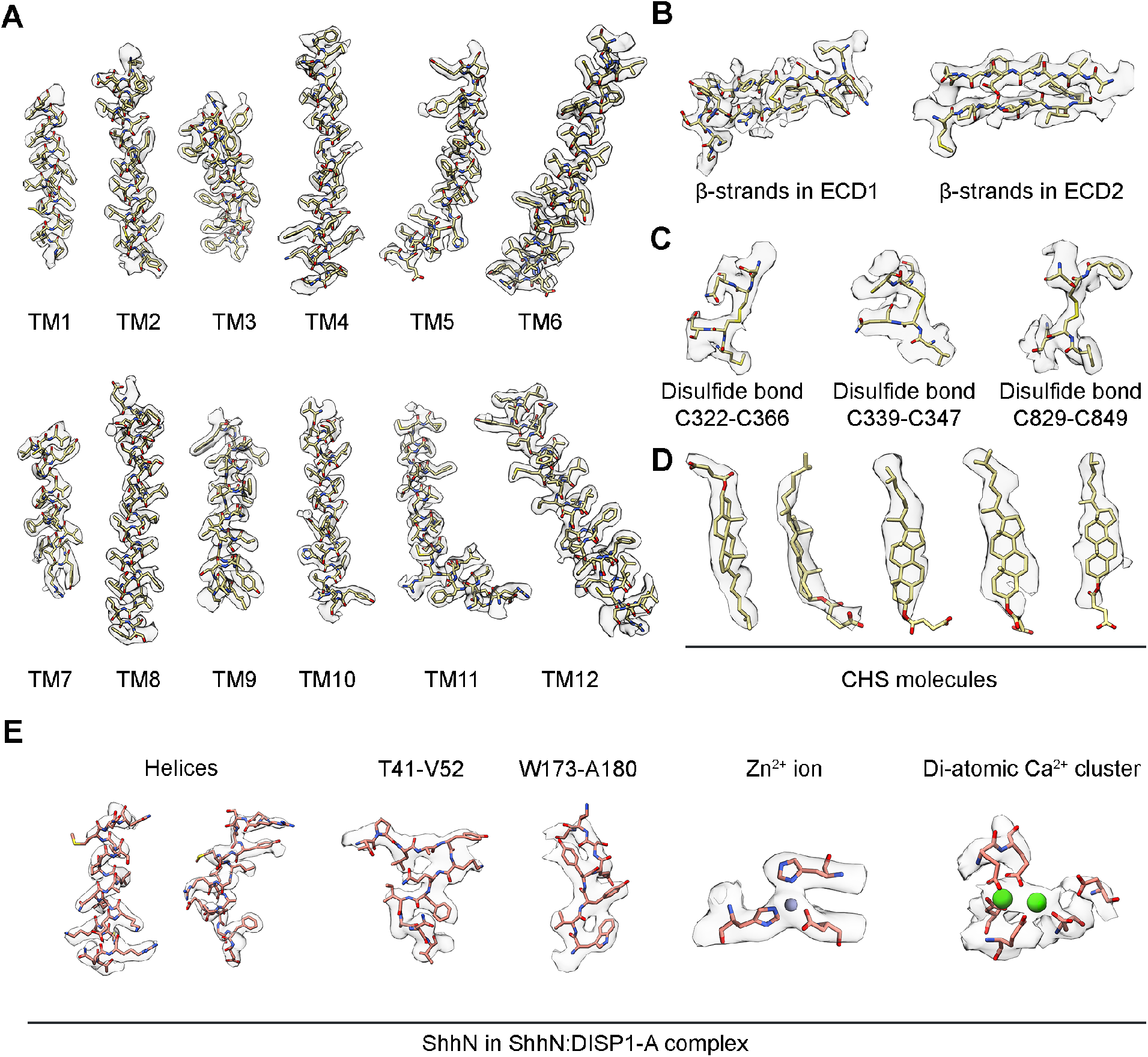
Representative cryo-EM densities from selected structural features. **A-D**, Representative cryo-EM densities from 3D reconstruction of DISP1-A, conformation R. **A**, Densities of all transmembrane helices; **B**, Representative densities of a beta-sheet from ECD1 and ECD2. **C**, Cryo-EM densities of three representative disulfide bonds. **D**, Cryo-EM densities of five representative CHS molecules. **E**, Representative cryo-EM densities from 3D reconstruction of ShhN:DISP1-A complex.

**Fig. S7.**
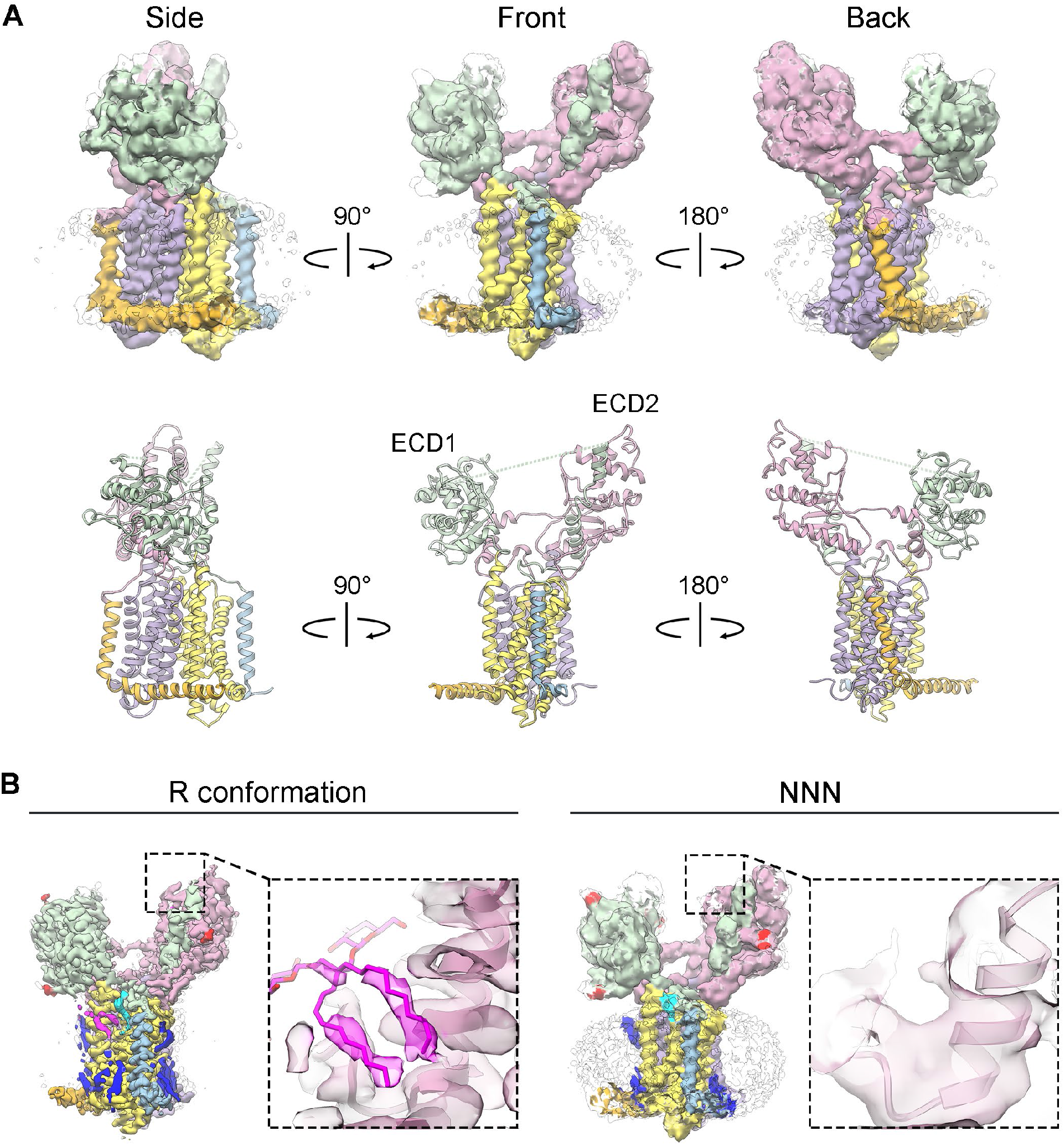
Structure and features of DISP1-A-NNN. **A**, Corresponding views of the DISP1-NNN cryo-EM density and atomic model are shown in the top and bottom rows; colors as in Fig. 1A,B. **B**, Comparison of cryo-EM densities in the region of the distal ECD2 cavity (small dashed box, enlarged to the right) for the R conformation of DISP1-A (5.3 sigma contour) and DISP1-A-NNN (3.4 sigma contour). Density is displayed as a pale line drawing, with the cavity occupied by LMNG (magenta) in DISP1-A. Note the absence of density within the DISP1-A-NNN cavity, even at a density threshold showing the entire micelle.

**Fig. S8.**
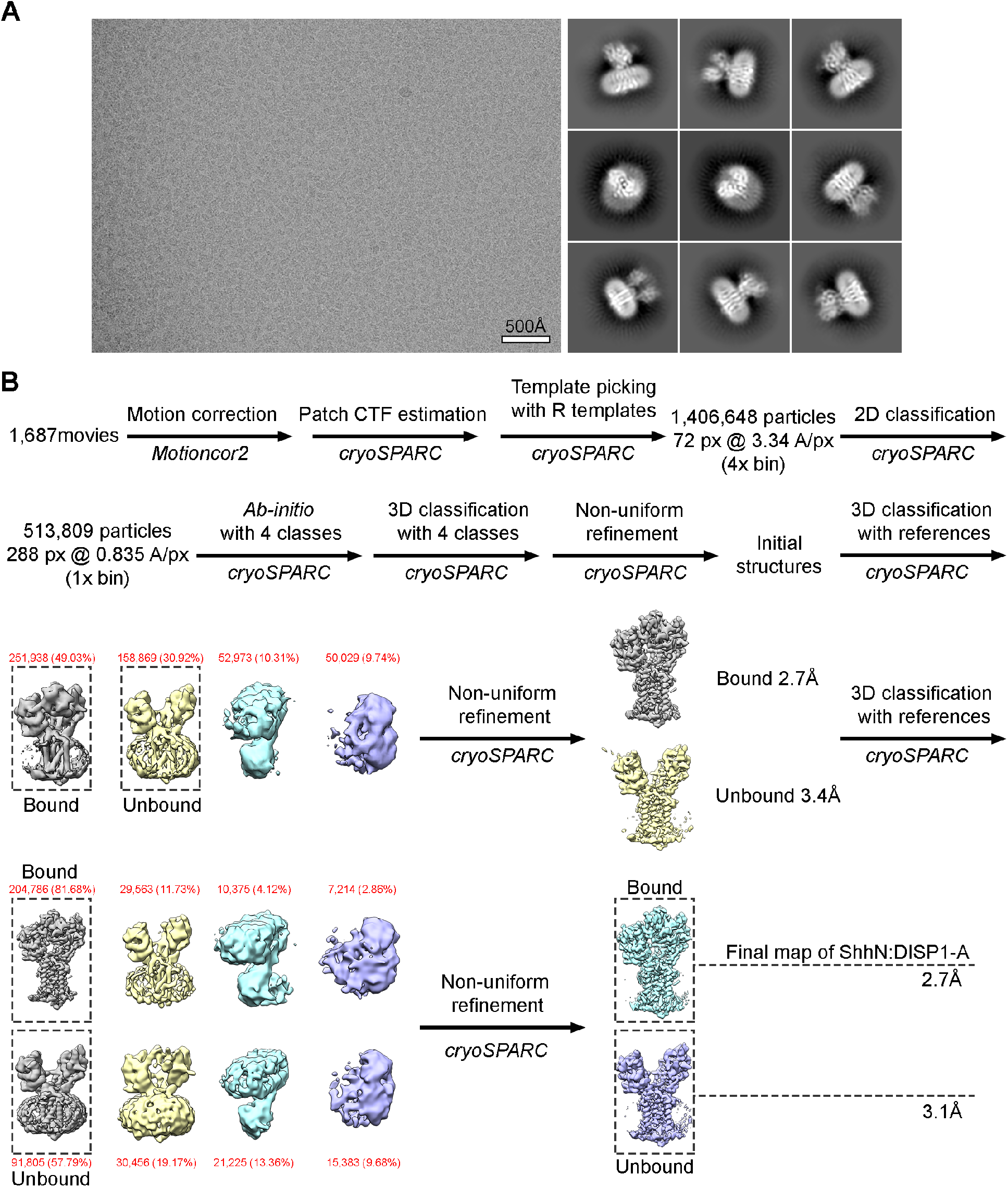
Cryo-EM data and image processing flow for ShhN:DISP1-A complex. **A**, A representative cryo-EM micrograph and highly populated reference-free 2D class averages for ShhN:DISP1-A. **B**, Schematic flow-chart illustrating the image processing used for ShhN:DISP1-A data. Thumbnail images are shown for reference-based 3D classes and high resolution refinements. Dashed black boxes indicate 3D classes selected for the next processing step, with class particle counts in red and refinement GS-FSC resolutions in black. The label, “3D classification with references,” indicates that explicit “apo” and “complex” references were used to seed the classification.

**Fig. S9.**
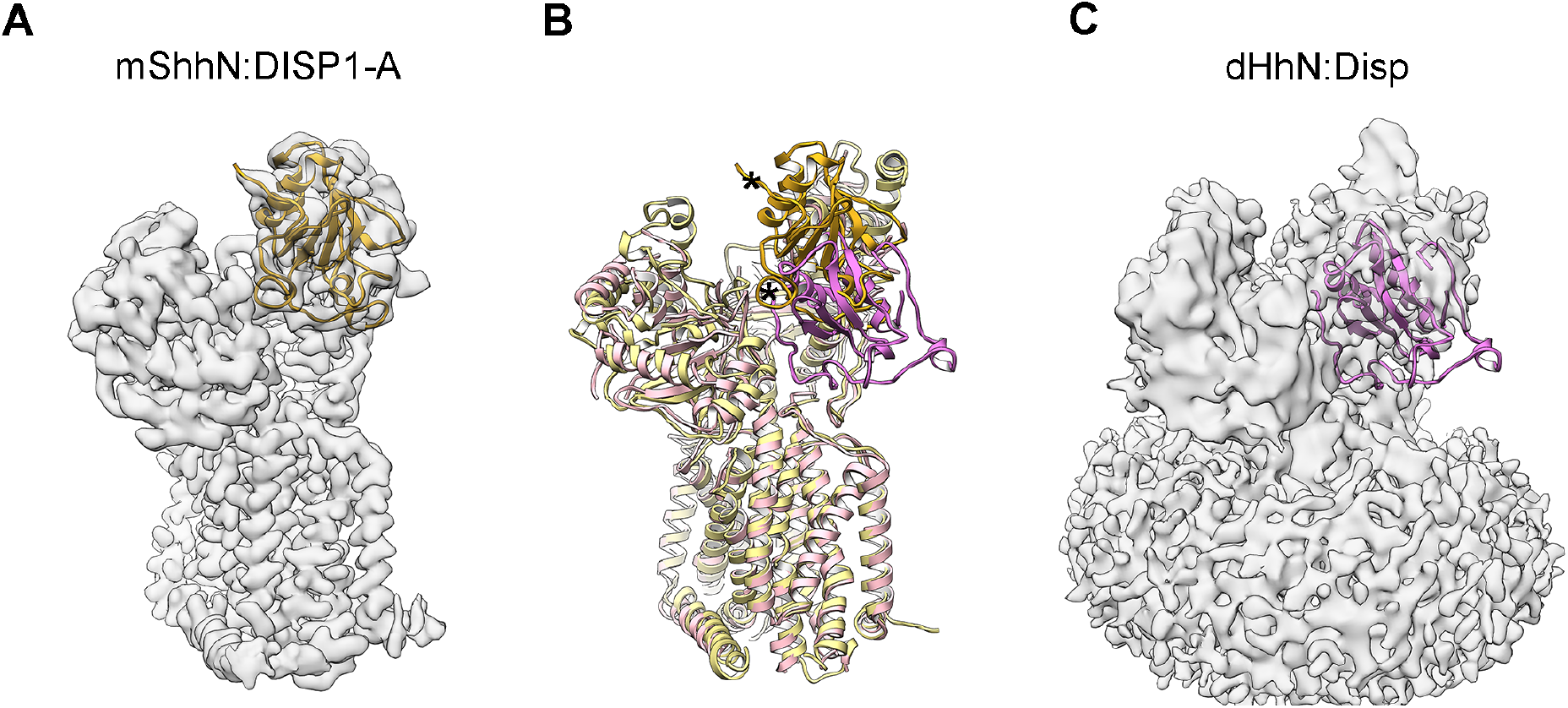
Relative positions assigned to murine and *Drosophila* Hedgehog proteins in complex with Dispatched. **A**, The cryo-EM density from the ShhN:DISP1-A complex in the current work (transparent grey), overlaid on a ribbon diagram of ShhN (goldenrod). **C**, The cryo-EM density from the *Drosophila* HhN:Disp complex in Cannac et al.[33] (transparent grey, EMD: 10464), overlaid on the ribbon diagram of HhN (orchid, extracted from PDB:6td6). **B**, Comparison of the ShhN:DISP1-A and HhN:Disp complex models reported here and in Cannac et al.[33]. Mouse DISP1-A and ShhN are in khaki and goldenrod, respectively, whereas *Drosophila* Disp and HhN are in pink and orchid. Relative to the *Drosophila* HhN model, murine ShhN is translated upwards, away from the membrane, and rotated towards the right. The asterisks indicate corresponding positions near the amino-termini of ShhN and HhN proteins.

**Fig. S10.**
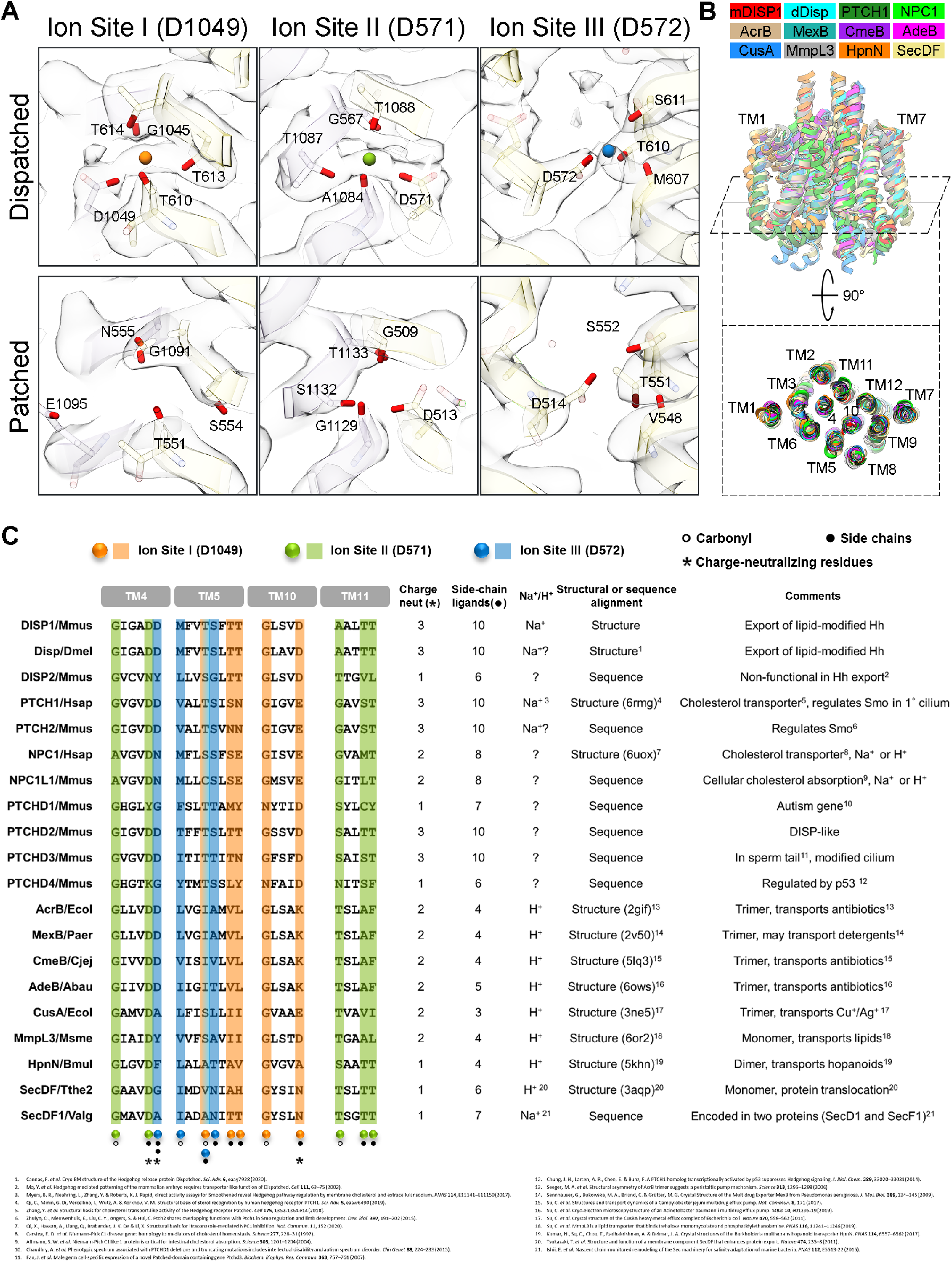
Na^+^ ion utilization by DISP1, PTCH1, and other members of the RND transporter family. **A**, Close-up views of Na^+^ ion sites I, II, and III in DISP1, showing the locations of liganding oxygens from amino acid side chains and main-chain carbonyls (see main text), and of the corresponding locations in PTCH1 (PDB ID: 6rmg), based on structural alignment of the two proteins. Note the presence in both proteins of a charge-neutralizing acidic residue at each site, and the conservation of side-chain oxygens as ligands. **B**, Structural alignment of RND family members, including DISP1, PTCH1, NPC1, and several prokaryotic RND transporters. **C**, Tabulated conservation of charge-neutralizing residues (3 total) or oxygen ligands from amino acid side-chains (10 total) in the indicated proteins, aligned from structure (**B**) or, if possible without ambiguity, aligned from sequence. Note the close conservation of Na^+^-liganding side-chains and charge-neutralizing residues in PTCH1, known to require Na^+^ for its activity, and in Disp from *Drosophila melanogaster*. The prokaryotic Na^+^-utilizing SecD1/SecF1 peptide translocator from *Vibrio alginolyticus*(encoded as two peptides; aligned by homology to *Thermus thermophilus* SecDF), in contrast, appears to have evolved a distinct mode of Na^+^ interaction. Abbreviations: Mmus, *Mus musculus*; Dmel, *Drosophila melanogaster;* Hsap, *Homo sapiens;* Ecol, *Escherichia coli;* Paer, *Pseudomonas aeruginosa;* Cjej, *Campylobacter jejuni;* Abau, *Acinetobacter baumannii;* Msme, *Mycolicibacterium smegmatis;* Bmul, *Burkholderia multivorans;* Tthe2, *Thermus thermophilus*; Valg, *Vibrio alginolyticus*.

**Table S1.**
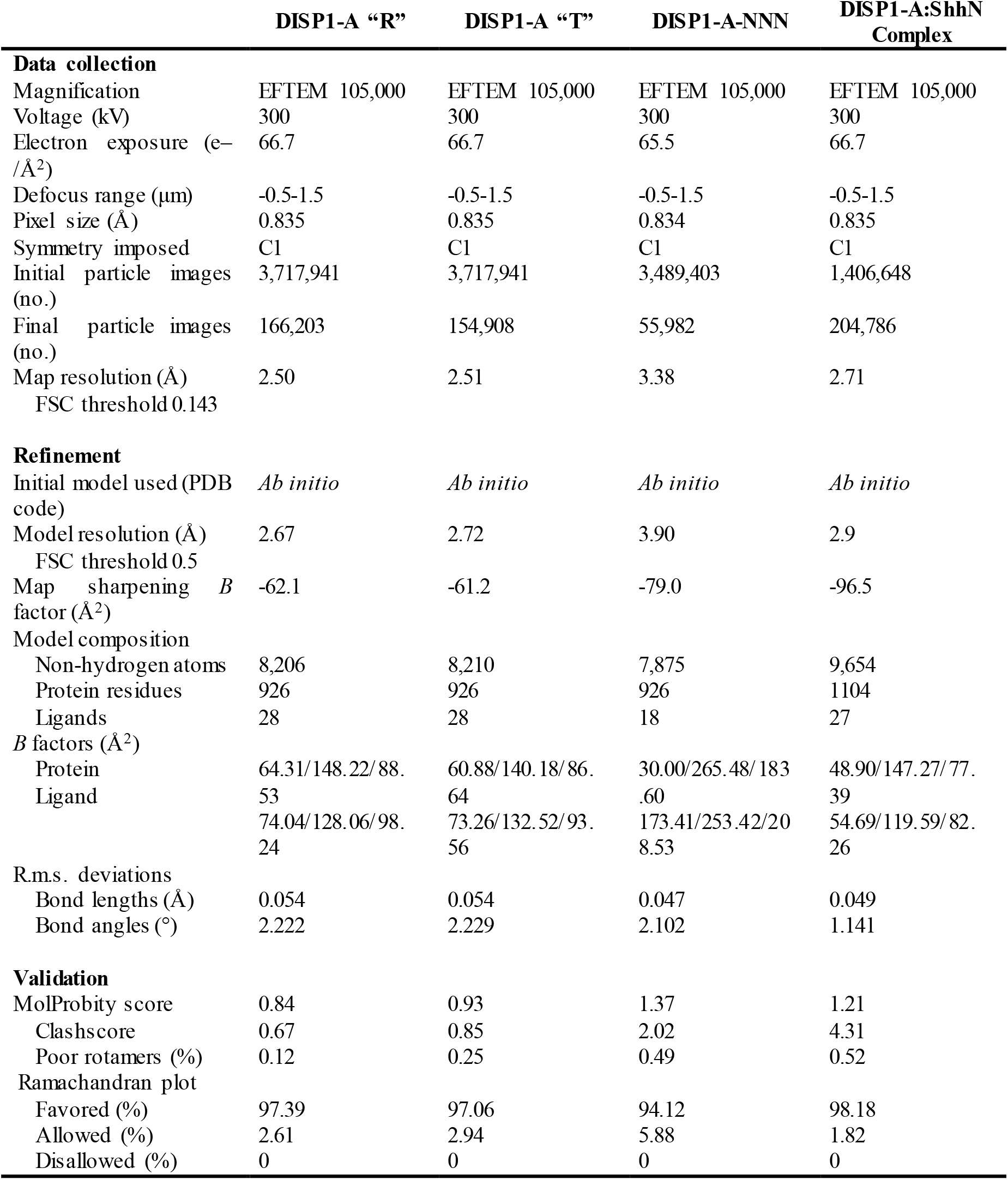
Cryo-EM data collection and model statistics.

